# Context-dependent effects of ketamine on emotional evidence integration

**DOI:** 10.64898/2026.10.30.762604

**Authors:** Miruna Rascu, Jan Grohn, Sara Costi, Caroline Zangani, Samuel Agyapong, Katharine A. Smith, Philip J. Cowen, Laurence T. Hunt, Miriam C. Klein-Flügge, Lilian A. Weber

## Abstract

Understanding others’ emotions requires continuously integrating noisy affective information as emotional states change over time. Research on emotion processing, however, has thus far focused on static environments or discrete events. Here, we developed a continuous emotional evidence integration task in which participants had to track the average emotional state in a rapidly changing sequence of faces under different levels of environmental volatility. This allowed us to examine emotion integration in changing environments. In online (n=92) and in-person (n=29) cohorts, participants integrated emotional information over longer timescales in stable environments while relying more strongly on recent evidence under volatility, consistent with principles established in perceptual decision-making. We further examined whether this process was modulated by ketamine, an NMDA receptor antagonist associated with evidence integration in the perceptual domain. We conducted a double-blind, placebo-controlled, within-subject study and probed emotion integration during the post-acute period, approximately 1.5 hours after the infusion. Ketamine modulated the temporal weighting of emotional evidence, shifting the balance between recent and past information depending on environmental volatility. Together, these findings show that emotional evidence integration dynamically adapts to the statistical structure of the environment and that ketamine can further modulate this process, which may have relevance for affective disorders.

## Introduction

In daily social interactions, humans continuously monitor others’ emotional expressions, integrating affective cues to infer underlying emotional states. During a conversation, for example, subtle shifts in another person’s facial expression can signal changes in their underlying emotional state and prompt an adjustment in behaviour. Such inferences depend not only on noisy incoming sensory information, but also on prior expectations about how stable or volatile others’ emotions are over time. A large body of work has examined how humans perceive and categorise emotional expressions, identifying key behavioural and neural mechanisms underlying emotion recognition (Adolphs, 2002; Barrett, 2017; Ekman, 1992; Lindquist et al., 2012). However, this literature has relied predominantly on discrete, trial-based paradigms using static stimuli. Although recent work has begun to examine how affective cues are integrated in dynamic scenes (Chen & Whitney, 2019; Ortega et al., 2025) we still have limited insight into how emotional information is integrated over time in changing and uncertain environments, where affective states unfold continuously.

By contrast, in the perceptual domain, extensive work has characterised how humans integrate continuously changing, noisy sensory information to guide adaptive behaviour. For example, we understand how humans accumulate and weight noisy evidence across sensory modalities (Brunton et al., 2013; Gold & Shadlen, 2007; Keung et al., 2019, 2020; Loosen et al., 2024; Wyart et al., 2012) and how these processes adapt to environmental statistics such as volatility and noise (Foucault et al., 2025; Glaze et al., 2015; Iglesias et al., 2013; Krishnamurthy et al., 2017; Mikus et al., 2025; Nassar et al., 2010, 2012; Ossmy et al., 2013; Ruesseler et al., 2023; Weber et al., 2020). In volatile environments, where underlying states change frequently, adaptive behaviour requires greater weighting of recent information, whereas stable contexts favour longer-term integration of past evidence (Behrens et al., 2007; Kilpatrick et al., 2019; C. Mathys et al., 2011; C. D. Mathys et al., 2014; Veliz-Cuba et al., 2016). Disruptions in volatility estimations have also been linked to psychiatric conditions such as schizophrenia (Cole et al., 2020; Deserno et al., 2020; Mikus et al., 2025), anxiety (Browning et al., 2015), autism (Lawson et al., 2017), and OCD (Vaghi et al., 2017). It is unclear, however, whether these principles of evidence accumulation and adaptive weighting under volatility guiding perceptual decisions generalise to domains such as emotional inference, where information is similarly noisy and unfolds over time. This is the first question addressed in this study.

Second, we consider the neural mechanisms that might underlie evidence integration in the emotional domain. In perceptual inference, N-methyl-D-aspartate (NMDA) receptors are required for the integration of noisy sensory information and the flexible updating of beliefs under uncertainty (Cavanagh et al., 2020; Corlett et al., 2006; Corlett, 2017; Corlett et al., 2019; Salvador et al., 2022; Wong & Wang, 2006). Clinical populations with glutamatergic dysfunction show altered volatility estimation and belief updating (Deserno et al., 2020; C. M. Kaplan et al., 2016; Mikus et al., 2025). Ketamine, a non-competitive NMDA receptor antagonist with known psychotomimetic effects (Corlett et al., 2006; Krystal et al., 1994, 2003; Moghaddam et al., 1997), has been shown to alter perceptual inference (Corlett et al., 2011; Mohanta et al., 2021; Weber et al., 2020; Weilnhammer et al., 2025) and uncertainty estimation (Cavanagh et al., 2020; Salvador et al., 2022; Vinckier et al., 2016) at subanaesthetic doses, consistent with disruptions in predictive processing and belief updating. These cognitive effects on perceptual evidence integration have predominantly been characterised during acute ketamine exposure.

In a distinct clinical literature, ketamine has been shown to elicit rapid and sustained improvements in depressive symptoms that emerge and persist beyond the acute drug phase (Abdallah et al., 2015; Berman et al., 2000; Vollenweider & Kometer, 2010; Zarate et al., 2006). Whether these psychotomimetic and antidepressant effects might share a common underlying mechanism remains unclear. Within a predictive coding framework (Friston, 2010; Friston & Kiebel, 2009), NMDA receptor-mediated signalling has been linked to regulating the precision of prior beliefs and determining the relative weighting of past expectations and incoming evidence (Corlett, 2017; Corlett et al., 2009; Sterzer et al., 2018). A reduction in prior precision may provide a potential common mechanism for both ketamine’s psychotomimetic and rapid antidepressant effects (Haarsma et al., 2021). In healthy individuals, it may lead to more flexible but less constrained inference, contributing to the psychotomimetic experiences, whereas in depression it may weaken overly precise and rigid negative priors, such as negative affective biases (Costi et al., 2025; Hinchcliffe et al., 2024, 2025; Pulcu et al., 2022; Stuart et al., 2015), increasing cognitive flexibility and allowing new, more positive information to carry greater weight in belief updating (Bottemanne et al., 2022).

Ketamine has also been shown to directly modulate emotional perception, with studies reporting changes in behavioural performance and neural responses to emotional faces (Lundin et al., 2021; Nugent et al., 2019; Reed et al., 2018, 2019). Moreover, recent computational work suggests that mood itself emerges from the ongoing integration of affective information, shaped by the relative weighting of prior expectations and new evidence (Asutay et al., 2021; Dercon et al., 2025; Eldar et al., 2016; Rutledge et al., 2014). If mood reflects the accumulation of affective information over time, then examining how ketamine modulates the integration of affective information may provide insight into the mechanisms underlying its rapid antidepressant effects. Our second question was therefore whether ketamine influences the integration of emotional evidence in the post-acute phase, when its antidepressant effects begin to emerge.

Here, we address these questions by combining approaches from perceptual decision-making with emotion research to characterise how affective information is continuously integrated over time, and to ask how this process is altered by ketamine. We developed a novel task in which noisy emotional stimuli, varying in valence or arousal, are presented in rapid succession, requiring participants to accumulate evidence to track the underlying affective state. This approach provides a more naturalistic framework for quantifying emotional evidence integration and allows us to examine how such processes adapt to environments with different levels of volatility. In a double-blind, placebo-controlled, within-subject design, we assessed continuous emotional evidence integration following ketamine or placebo administration. Behaviour was measured at a post-acute time point (1.5 hours following the end of the drug infusion), when transient psychotomimetic effects have subsided, given ketamine’s rapid distribution (*α* ≈ 7 minutes) and elimination (*β* ≈ 2-4 hours) half-lives. This timepoint was chosen to capture the early changes in affective and cognitive processing potentially related to ketamine’s antidepressant effects reported in clinical populations (Zarate et al., 2006).

In preregistered analyses (https://aspredicted.org/bp4w-9t7p.pdf), we find that participants track dynamically changing emotional signals, show emotion-specific biases, and adapt the temporal weighting of emotional evidence to environmental volatility when rating valence, most clearly in a large online sample. Contrary to our prediction, ketamine did not increase reliance on recent evidence overall. Instead, it modulated the volatility dependence of emotional evidence weighting. Following ketamine, but not placebo, recent evidence was weighted more strongly in volatile than in stable environments, in the in-person sample. These findings suggest that emotional processing may be better understood as a dynamic process of evidence integration than as the evaluation of isolated stimuli, opening a new route to understanding how mood disorders, and rapid-acting antidepressants such as ketamine, shape the way humans update emotional beliefs.

## Results

### Participants reliably tracked changes in valence and arousal levels

We designed a novel task (**Fig 1**) in which participants were asked to continuously integrate noisy emotional evidence to track a dynamically changing underlying true mean level of emotion. Two rating dimensions were tested in separate blocks: participants tracked either the mean level of valence (very negative to very positive) or arousal (low to high emotional intensity). A rapidly changing continuous stream of faces expressing different emotions was presented in each block, with individual face stimuli sampled from a normal distribution centred around a latent true mean. This underlying true mean (**Fig 1, second row**, thick blue and pink lines) occasionally changed within a block (change-points), requiring participants to update their estimates by observing and integrating information across successive samples. Ratings could be updated at any time, but no feedback was provided during the block. Individual face stimuli varied in emotional valence and arousal on a continuous scale from approximately 1-8, based on normative ratings provided with the KDEF-dyn database (Calvo et al., 2018; Lundqvist et al., 1998). The underlying true mean valence and arousal took values 2-6 for valence and 2, 4, and 6 for arousal. Participants reported their estimates of the underlying true mean using a five-point scale for valence (with each point on the scale corresponding to one possible true mean value) and a three-point scale for arousal to simplify arousal judgments, which were found to be more difficult during task piloting. Their task was to track the average emotional response of an audience, inferred from the stream of facial expressions, while the audience watched different “movies” which varied in how rapidly they elicited changes in emotion (Movie A vs Movie B). The two movies represented different types of environments, balanced across valence and arousal: blocks with low volatility (less frequent changes in the true underlying mean, 16 change-points on average per block) and blocks with high volatility (around 31 change-points on average per block). Participants learned these different environmental statistics during training, without being explicitly told which “movie” was more volatile. Change-points also varied in magnitude: valence shifts could be small (1-step, corresponding to one scale step), medium (2-step), or large (3-step), whereas arousal blocks included small (2-step, corresponding to one scale step) and large (4-step, two scale steps) changes. Given the different response scales and change-point magnitude levels across rating dimensions, valence and arousal are considered separately across analyses.

**Figure 1.**
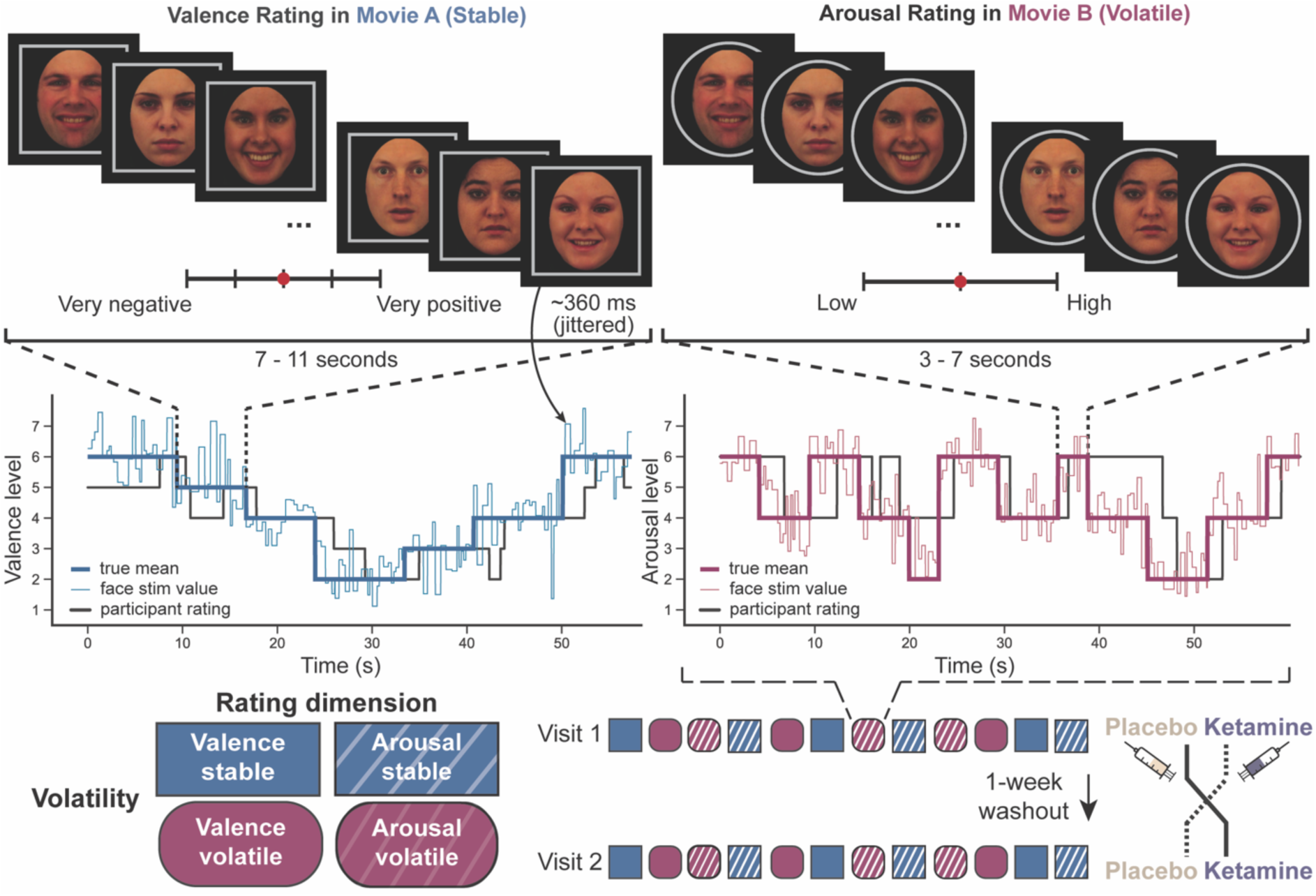
Continuous emotion rating task structure. Participants viewed a rapid stream of facial stimuli and continuously rated either the average valence (from very negative to very positive; 5-point scale) or arousal (low to high emotional intensity; 3-point scale), in separate blocks, moving a slider with left/right button presses. **Top:** example task displays from a valence-rating block in the stable environment (Movie A; left) and an arousal-rating block in the volatile environment (Movie B; right). Frame shapes around the face served as cues indicating the ‘movie’, i.e. the statistical properties of the environment, with square frame for stable and circular frame for volatile conditions. At the start of each block, participants were informed of the rating dimension (valence or arousal) and movie type (A or B). **Middle:** example stimulus sequences for a stable valence block (left) and a volatile arousal block (right). Thick lines show the underlying true mean (blue, stable; pink, volatile), which changed every 7-11 s in stable and every 3-7 s in volatile blocks. Thin lines indicate the valence or arousal value of individual face stimuli, and black lines show a representative participant’s ratings. **Bottom left:** the four task conditions (rating dimension x volatility). Colour and shape indicate volatility (blue rectangle, stable; pink circle, volatile), and fill pattern indicates rating dimension (solid, valence; striped, arousal). **Bottom right:** task conditions were counterbalanced across the session, with three blocks of 2.5 minutes per condition. In-person participants completed the task in two visits, separated by a 1-week washout, that were identical except for the intravenous infusion of either ketamine or placebo (counterbalanced). The block sequences were identical across the two visits. Faces shown are KDEF-dyn stimuli F09-NE-HA, F03-NE-DI, M10-NE-SU, F29-NE-HA, F20-NE-HA and M12-NE-HA (Calvo et al., 2018; Lundqvist et al., 1998).

The task was completed by two independent samples: an online cohort (n = 92) and an in-person cohort (n = 29). The in-person participants performed the task in two study visits, one week apart, approximately 1.5 hours after the end of the ketamine or placebo infusion. Results reported in the main text are based on the in-person cohort, and those unrelated to drug infusion were replicated in the independent online sample (**Fig S1, Supplementary Tables S5-S8**), unless stated otherwise. Where the cohorts differed, both results are reported. To validate the task design, we first assessed whether participants successfully tracked the dynamically changing emotional mean. As a visual illustration of response dynamics, we examined the cumulative proportion of detected change-points over time following change-point onset (Kaplan-Meier estimate; see Methods), where a change-point was considered detected once participants changed their rating in the same direction as the underlying shift in emotional mean (**Fig 2A**). Detection began to rise steeply approximately 1 second after the change-point onset, with half of all change-points detected within approximately 2.5s and continued to increase more slowly afterwards. This time course is consistent with participants accumulating evidence across several faces before adjusting their rating. Arousal showed a comparable overall time course of response but with slightly lower detection than for valence over time.

**Figure 2.**
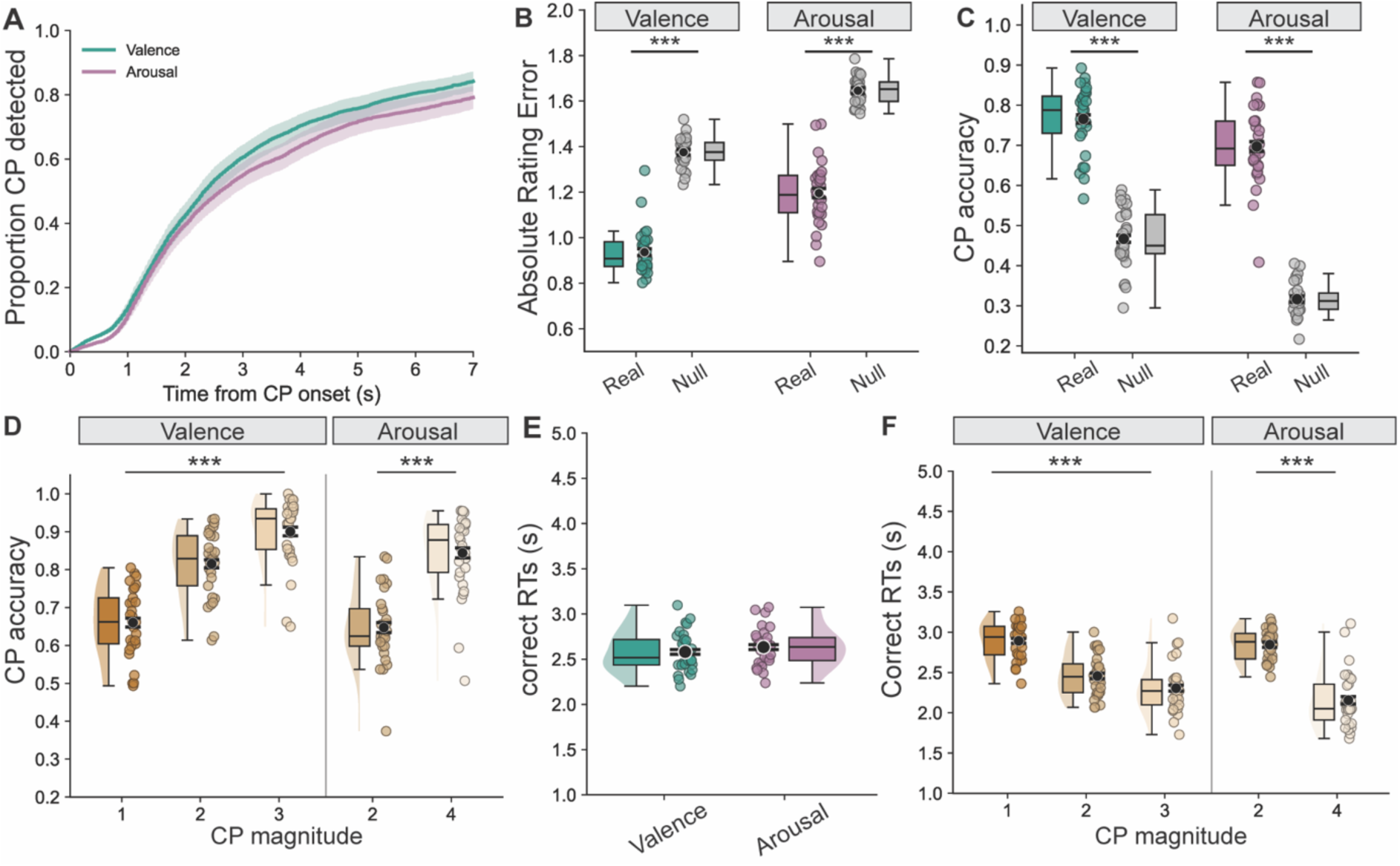
Behavioural performance for rating valence and arousal during continuous emotional evidence integration. **A.** Cumulative proportion of detected change-points as a function of time from change-point onset, shown separately for valence and arousal (Kaplan-Meier estimate, mean ± 95% CI). A change-point was considered detected once participants made a rating change in the same direction as the underlying mean shift. Responses were counted from change-point onset and change-points that ended before a response were treated as censored (see Methods). Arousal showed slightly lower detection accuracy than valence. **B.** Continuous tracking performance, as absolute rating error (|participant rating - true underlying mean|). Rating errors were substantially smaller in real data than in shuffled data. **C.** Change-point (CP) detection accuracy exceeded a frame-shuffled null model in which responses were permuted within each block while preserving rating durations, confirming that participants used available emotional information rather than responding randomly. **D**. Change-point detection accuracy increased with CP magnitude (the size of the change in the true mean) for both valence and arousal, indicating increased sensitivity to stronger changes in evidence. **E.** Reaction times (RTs) were similar in valence and arousal. **F.** RTs decreased with larger CP magnitudes for both valence and arousal, showing faster responses when the underlying mean shift was more pronounced. Black dots and error bars denote group level means ± within-subject s.e.m. (Cousineau-Morey correction), whereas coloured dots denote participant level data. *** p < .001.

To examine whether people were able to track the underlying mean, we quantified performance using two complementary measures. First, we computed a continuous, framewise metric based on the absolute deviation between the participant’s rating and the true underlying mean across the whole block (rating error). This measure captured how accurately participants tracked the true mean over time, including during long stable periods when no change-points occurred. To ensure participants did not respond randomly we estimated a null distribution obtained by shuffling participants’ rating runs within each block (see Methods for details). Continuous absolute rating errors were markedly smaller in the real data than in shuffled data (valence: real M = 0.94 vs null M = 1.38, paired t-test: t(28) = -22.61, p < .001, d = -4.20; arousal: real M = 1.20 vs null M = 1.65, t(28) = -16.83, p < .001, d = -3.12; **Fig 2B**; see Methods for details), confirming that participants’ ratings tracked meaningful structure in the emotional stimulus stream.

Second, we characterised responses to change-points (i.e. shifts in the true underlying mean). Using the same directional-update criterion as above, we computed change-point based accuracy, defined as whether participants adjusted their rating in the correct direction within a predefined detection window (see Methods for details). A response was considered correct if the rating change within this window matched the direction of the shift in the true underlying mean. We then quantified response times (RTs) for correctly detected change-points. These change-point-based metrics, commonly used in perceptual change-point detection tasks (Booras et al., 2021), allowed us to test whether participants were sensitive to discrete shifts in the underlying emotional mean. Participants reliably detected change-points, performing well above chance, as estimated using the null model (valence: real M = 0.76 vs null M = 0.47, t(28) = 28.87, p < .001, d = 5.36; arousal: real M = 0.70 vs null M = 0.32, t(28) = 31.51, p < .001, d = 5.85; **Fig 2C**). We then tested whether performance scaled with the size of the change in the true underlying mean. Indeed, sensitivity to changes increased with signal strength, i.e. the magnitude of the underlying mean shift: larger change-points yielded higher change-point accuracy in both valence (main effect of change-point magnitude: b = 0.888, 95% CI [0.763, 1.014], χ²(1) = 61.49, p < .001, **Fig 2D**) and arousal (b = 0.610, 95% CI [0.511, 0.709], χ²(1) = 55.60, p < .001).

Having characterized participants’ response accuracy, we turned to RTs to see if these provided further insight into people’s emotion integration performance. RTs were relatively slow overall (global M = 2.60 s, SD = 0.19), consistent with participants integrating evidence over multiple facial stimuli before adjusting their rating. We found no evidence that RTs differed between valence and arousal (arousal: M = 2.63 s, SD = 0.21; valence: M = 2.58 s, SD = 0.22; main effect of rating dimension on logRT: b = 0.011, 95% CI [-0.017, 0.039], F(1, 27.9) = 0.63, p = .434; **Fig 2E**). Analogous to effects seen for CP accuracy, RTs again decreased with change-point magnitude for both dimensions (main effect of change-point magnitude, valence: b = -0.110, 95% CI [-0.130, -0.090], F(1, 27.9) = 125.18, p < .001; arousal: b = -0.136, 95% CI [-0.152, -0.120], F(1, 29.2) = 315.86, p < .001, **Fig 2F**), consistent with faster responses to more salient changes in the underlying emotional signal.

### Emotion-dependent biases and prioritization in continuous emotional evidence integration

Although participants successfully tracked the underlying emotional mean, emotional information is unlikely to be processed as an objective signal and may introduce systematic biases into evidence integration. For example, people may detect and integrate faces with negative emotion more rapidly compared to positive emotion, and similarly for faces with high arousal compared to low arousal. We therefore examined whether tracking performance was influenced by the direction of change in the true underlying mean (upwards or downwards) and the emotional level reached (valence: negative, neutral and positive; arousal: low, medium, high). For valence, we separately considered mean shifts towards negative (true mean valence levels 2-3), neutral (level 4) and positive (levels 5-6) emotion levels, whereas in arousal we considered mean shifts towards low (true mean arousal level 2), medium (level 4) and high (level 6).

We first assessed whether overall tracking accuracy varied across emotion levels using the continuous signed rating error metric (participant rating – true mean; **Fig 3A**). To compare rating errors at opposite ends of each rating dimension, errors at the extreme levels (negative and positive valence, low and high arousal) were sign-flipped for positive valence and high arousal, so that larger values at either end indicate ratings that were less extreme than the true mean, i.e. biased towards the centre of the scale. Errors at intermediate levels (neutral valence and medium arousal) were not flipped and are analysed separately below. When rating valence, participants had higher rating errors for positive compared to negative levels (main effect of emotion level: b = 0.132, 95% CI [0.027, 0.237], F(1, 28) = 6.595, p = .016), whereas in arousal rating errors were higher in low compared to high arousal levels (b = -0.453, 95% CI [-0.643, -0.264], F(1, 28) = 23.977, p < .001; **Fig 3A, left**). We then tested whether participants showed systematic biases when tracking intermediate emotion levels (neutral valence and medium arousal), where signed errors are expected to be centred around zero, given that deviations can occur in both positive and negative directions. For neutral valence, participants tended to rate expressions as slightly more positive than the true mean, although the signed rating error did not significantly differ from zero (b = 0.059, 95% CI [−0.002, 0.120], F(1, 28) = 3.893, p = .058; **Fig 3A, top right**), whereas the online cohort showed a small bias in the opposite direction (online cohort, valence neutral RE against zero: b = -0.064, 95% CI [-0.113, -0.015], F(1, 91) = 6.77, p = .011; **Fig S1D**). Medium arousal levels showed a strong positive bias (b = 0.469, 95% CI [0.356, 0.582], F(1, 28) = 72.534, p < .001; **Fig 3A, bottom right**), suggesting participants overestimated the true arousal at medium levels. Together, these analyses revealed systematic emotion-dependent asymmetries in continuous ratings, with lower errors for negative than positive valence and for high than low arousal, alongside a tendency to overestimate intermediate emotion levels.

**Figure 3.**
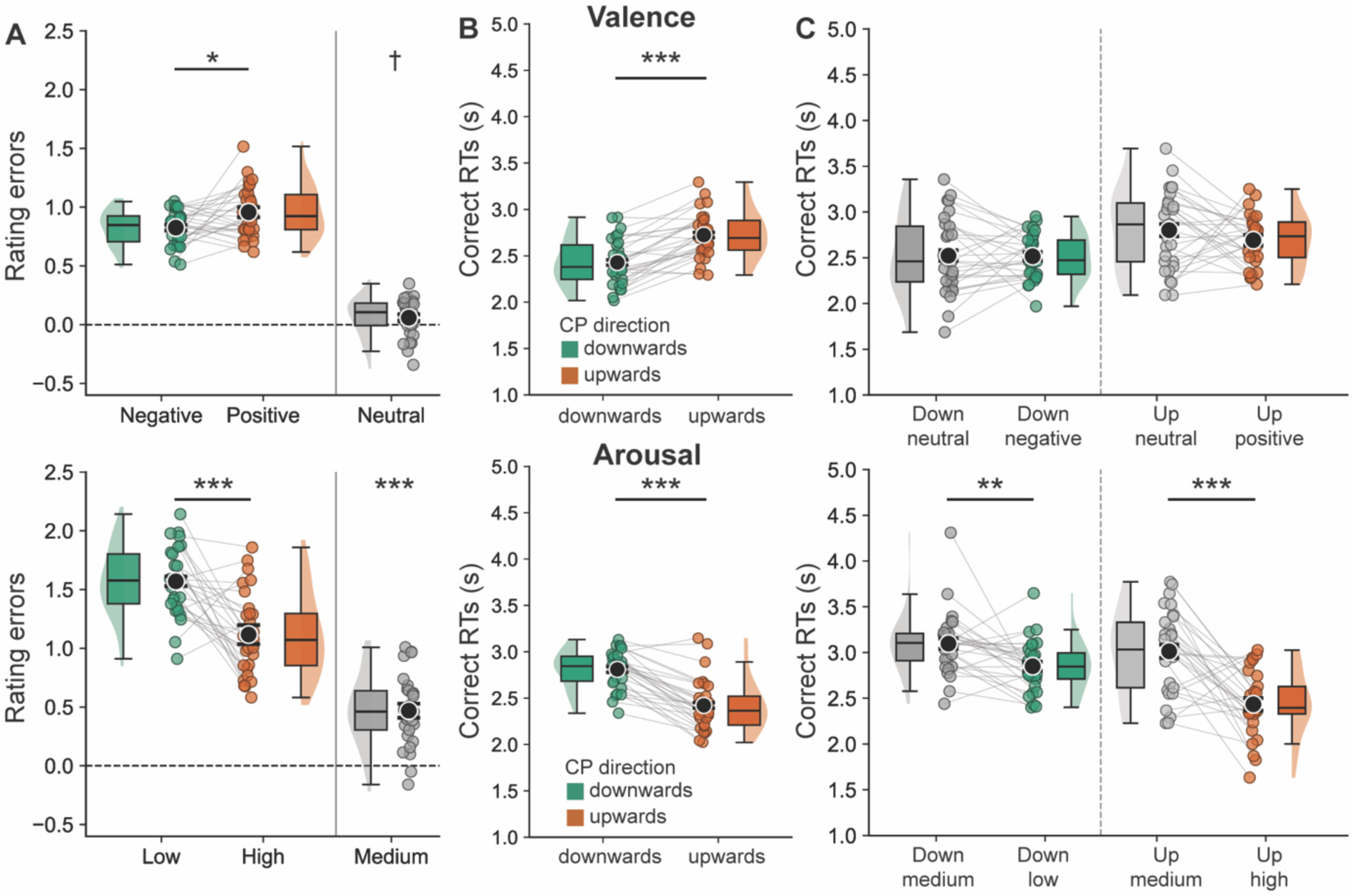
Distinct effects of change-point direction and emotion level during emotional evidence integration. Each row shows one emotional dimension (valence, top; arousal, bottom). **A.** Rating error (participant rating - true underlying mean) across emotion levels. For comparability, rating error for positive valence and high arousal was sign-flipped, so that larger values at both ends of the scale indicate ratings less extreme than the true mean (biased towards the centre of the rating scale). Rating errors were lower when the true mean shifted towards negative valence and towards high arousal. Rating errors for neutral valence and medium arousal are shown separately as signed errors. Because deviations around intermediate levels can occur in both directions, errors are expected to centre around zero. Medium arousal showed positive rating errors, indicating that participants tended to rate medium arousal levels as closer to high arousal. Neutral valence did not differ significantly from zero (p = .058) but showed a trend towards positive errors. **B.** Reaction times (RTs) for correctly detected change-points, grouped by direction of the shift in the true underlying mean (down vs up). RTs varied as a function of change-point direction, with faster responses for downward vs upward changes in valence, and for upward vs downward changes in arousal. **C.** Correct RTs grouped by change-point direction and the emotional level reached following the change. In valence blocks, RTs did not differ between the emotional levels reached (negative vs neutral for downward changes; neutral vs positive for upward changes), indicating that RTs were driven primarily by the direction of change, rather than the final valence level, prioritizing downwards changes towards more negative emotional valence. In arousal blocks, RTs differed between the emotional levels reached within both downward and upward changes (low vs medium for downward changes; medium vs high for upward changes), with slower responses towards medium than towards more extreme arousal levels. Black dots and error bars denote group level means ± s.e.m. with Cousineau-Morey correction, whereas coloured dots denote participant level data. *** p < .001, ** p < .01, * p <.05 and † p < .10.

We next turned to emotional biases in participants’ responses to emotional changes, asking whether sensitivity to changes in the true underlying mean depended on the direction of change (up towards positive valence or high arousal, or down towards negative valence or low arousal). In valence blocks, upward changes had longer RTs than downward changes when controlling for change point magnitude (main effect of CP direction: b = 0.091, 95% CI [0.063, 0.120], F(1, 27.9) = 43.81, p < .001, **Fig 3B, upper panel**), whereas in arousal, upward changes were associated with shorter RTs than downward changes (main effect of CP direction: b = -0.117, 95% CI [-0.148, -0.086], F(1, 29.8) = 59.37, p < .001, **Fig 3B, lower panel**). This is consistent with the rating error effect suggesting prioritised processing of changes towards more negative and towards higher arousal emotion levels. However, changes towards more negative emotions include changes from positive to neutral and from neutral to negative levels (and accordingly for arousal). To dissociate whether the observed effects were driven by the direction of the change or by the emotion level reached after the change, we split the analysis into CP direction and reached emotion level while controlling for CP magnitude. Given that reached emotion level and CP magnitude are not fully crossed, large CP magnitudes were excluded in this analysis (for all CPs see **Supplementary Table S2, Fig S2**). In valence blocks, RTs were not significantly different between emotion levels reached in either downward (neutral vs negative, main effect of reached emotion level: b = -0.004, 95% CI [-0.059, 0.050], F(1, 27.9) = 0.03, p = .867, **Fig 3C, upper panel**) or upward changes (positive vs neutral: b = -0.011, 95% CI [-0.073, 0.050], F(1, 27.6) = 0.14, p = .709). This indicates that in the in-person sample, valence RTs were primarily driven by the direction of change, rather than the final valence level. In the online sample, however, responses after downward changes were faster towards neutral than towards negative valence (b = -0.102, 95% CI [-0.160, -0.043], F(1, 111) = 11.87, p < .001). In arousal blocks, RTs significantly differed between emotion levels reached in both directions and both cohorts. RTs were slower when changing towards medium arousal levels (medium vs low levels: b = 0.086, 95% CI [0.036, 0.136], F(1, 27.5) = 12.27, p = .002; medium vs high levels: b = 0.214, 95% CI [0.137, 0.290], F(1, 27.7) = 32.46, p < .001; **Fig 3C, lower panel**), suggesting that more extreme arousal levels on either side were prioritized and responded to faster. Together, these results suggest distinct patterns of emotional biases in belief updating: valence biases are mainly direction-dependent, with a less consistent contribution of the level reached, whereas arousal biases are target-level dependent.

### Emotional evidence integration is sensitive to environmental volatility

Having established that participants were able to continuously track mean levels of emotional valence and arousal, and that certain emotions and arousal levels were prioritized in terms of accuracy and speed of responding, we next examined whether the environment volatility influenced participants’ rating behaviour. From a normative perspective, we predicted faster updating and greater reliance on recent evidence in volatile environments compared to more extended integration in stable ones. After controlling for change-point magnitude, direction and averaging across drug sessions, participants had faster reaction times to change-points in volatile than stable blocks when rating valence (main effect of volatility, valence: b = -0.084, 95% CI [-0.116, -0.051], F(1, 27.9) = 28.10, p < .001; **Fig 4A**) but not when rating arousal (main effect of volatility: b = 0.019, 95% CI [-0.021, 0.058], F(1, 27.9) = 0.96, p = .337). Faster RTs in volatile environments are consistent with more rapid updating in response to frequent changes. Additional analyses of rating change frequency and change-point accuracy also showed that behaviour was sensitive to environmental volatility (**Supplementary Results** and **Fig S3**).

**Figure 4.**
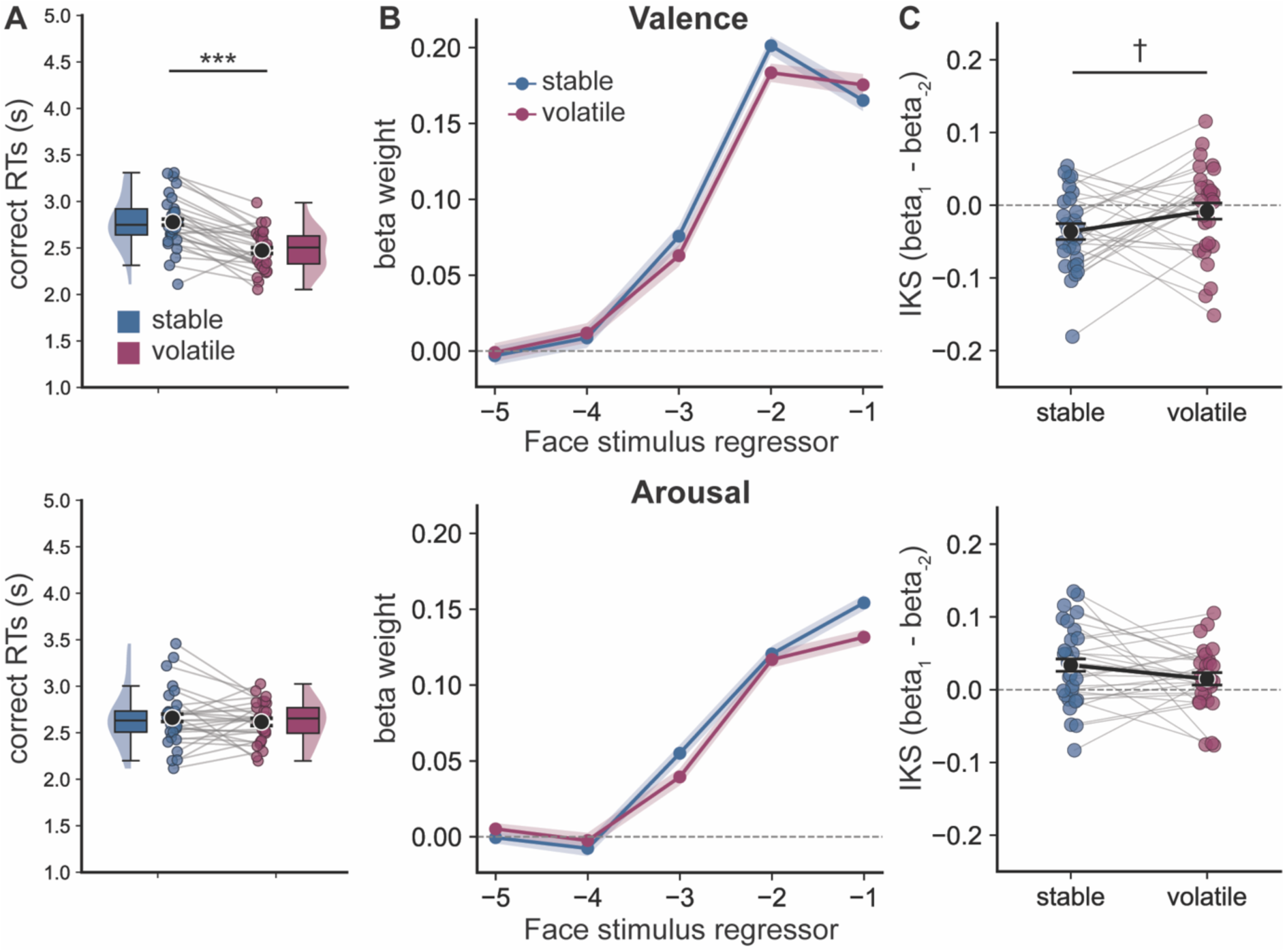
Participants adapt reaction times and emotional evidence integration kernels to environmental volatility. The task included two environmental conditions: stable blocks, where the underlying mean changed rarely, and volatile blocks, where the mean changed frequently. The top row shows blocks where participants rated valence, the bottom row arousal. **A.** Reaction times to detected change-points. Participants responded more quickly in volatile blocks than in stable blocks in valence but not in arousal, indicating that rapid environmental change elicited faster behavioural adjustments. **B.** Integration kernel beta weights for valence (top) and arousal (bottom). We estimated linear regression-based evidence-integration kernels by regressing participants’ rating updates onto the prediction errors (face value - current rating) of the five preceding faces. Weights were highest for the two most recent faces in both dimensions. Lines show mean kernel weights and shaded bands ± within-subject s.e.m. (Cousineau-Morey). **C.** Integration-kernel steepness (IKS) defined as the difference between the beta weight in the most recent face minus the weight in the second most recent face. Larger values indicate steeper, more recency-focused kernels. In person IKS did not differ significantly between volatility conditions in either dimension (valence p=.075, arousal p=.124). Black dots and error bars denote group level means ± s.e.m. with Cousineau-Morey correction, whereas coloured dots denote participant level data. *** p < .001, † p < .10.

To directly assess how participants adjusted the temporal weighting of evidence across conditions, we next examined integration kernels, which capture how strongly information from recent versus earlier stimuli influences current responses. We estimated linear regression-based integration kernels that quantify how strongly the prediction error of each of the five face stimuli preceding a rating update predicted the direction and size of that update (see Methods for details). This deviates from the preregistered (https://aspredicted.org/bp4w-9t7p.pdf) time-binned logistic regression, because the time-binned kernel depended on an arbitrary bin width and its binary outcome discards the direction and size of rating changes (see **Supplementary Deviations from preregistration;** the preregistered kernel is reported in **Supplementary Table S3b** and **Fig S4**). Across both valence and arousal, kernel weights were strongly biased toward recent evidence, with the largest contributions arising from the two most recent face stimuli and decaying weights for earlier faces (**Fig 4B)**, indicating a general recency-weighted integration strategy.

To quantify whether recency weighting changed with volatility, we used a preregistered summary metric of integration-kernel steepness (IKS), defined as the difference between the beta weights of the most recent and the second most recent face (**Fig 4C**). Larger IKS values indicate steeper, more recency-biased kernels. In valence, kernels were steeper in volatile than in stable blocks in the online cohort (online, main effect of volatility on valence IKS: b = 0.038, 95% CI [0.013, 0.062], F(1, 91) = 9.02, p = .003) but only at a trend level in the in-person sample, averaged across drug sessions (main effect of volatility on valence IKS: b = 0.028, 95% CI [-0.003, 0.059], F(1, 84) = 3.24, p = .075), consistent with an adaptive increase in recency weighting under higher volatility conditions. In arousal, there was no effect of volatility either online or in person (online main effect of volatility on arousal IKS: b = −0.005, 95% CI [−0.022, 0.012], F(1, 91) = 0.36, p = .550; in person: b = -0.019, 95% CI [-0.043, 0.005], F(1, 84) = 2.42, p = .124). Together, these results indicate that participants flexibly adjusted both the speed and temporal weighting of evidence integration in response to environmental volatility when rating valence but not arousal.

### Ketamine selectively modulates emotional evidence integration depending on volatility conditions

We next investigated how emotional processing and emotional evidence integration were modulated by ketamine in the in-person sample (n = 29, **Fig 5A**). The continuous emotion rating task was performed 1.5 hours after the end of the drug infusion, by which time the acute psychotomimetic and dissociative effects measured using the BPRS and CADSS-6 had largely resolved (**Fig S6A-B**).

**Figure 5.**
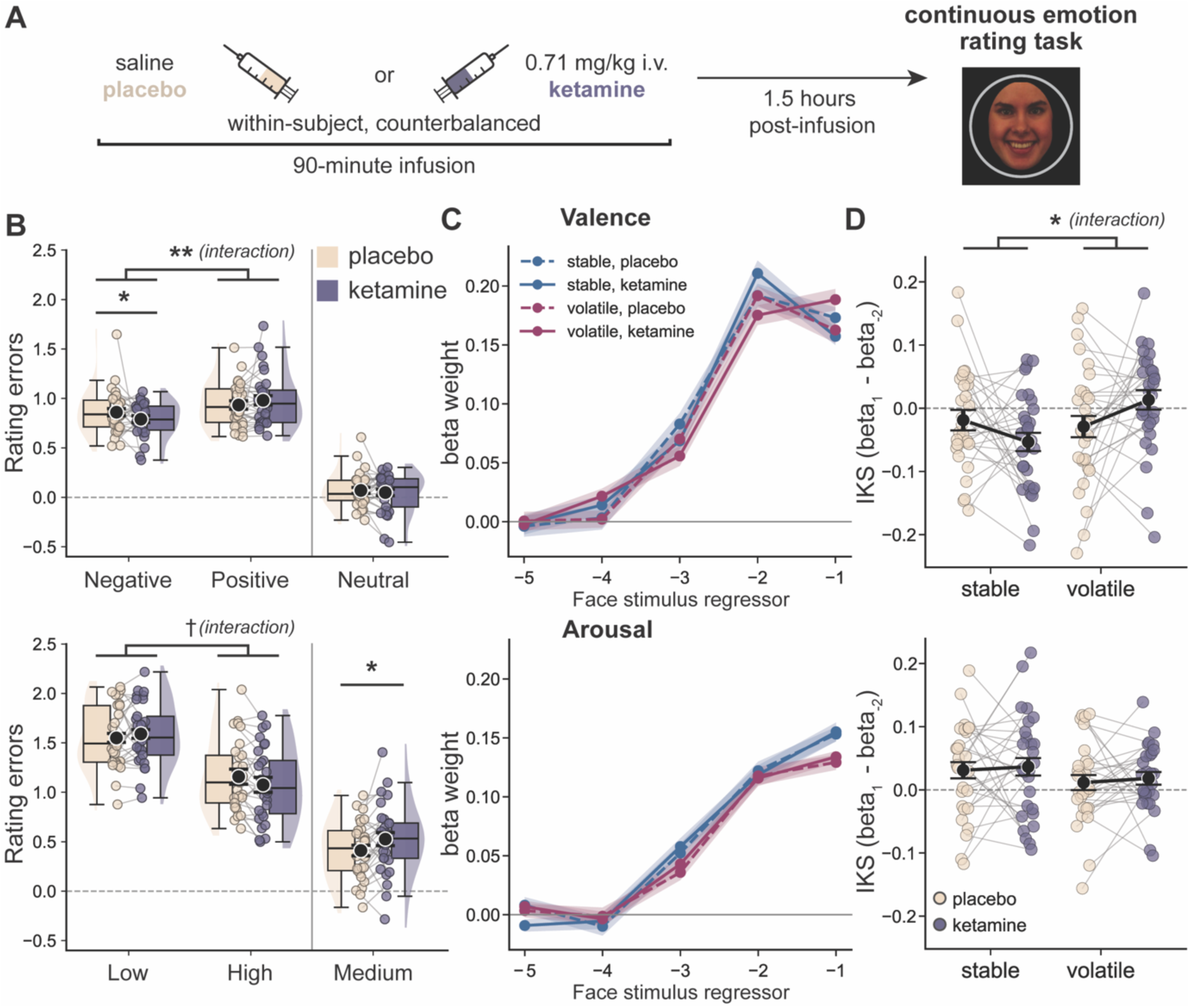
Ketamine produces small, volatility-dependent changes in the temporal integration of emotional evidence. **A.** Participants completed the continuous emotion rating task 1.5 hours following the end of the infusion of either ketamine or placebo in a within-subject, counterbalanced design across two sessions, one week apart. The face shown is from the KDEF-dyn stimulus F29-NE-HA (Calvo et al., 2018; Lundqvist et al., 1998). **B.** Ketamine modulated rating errors across negative and positive valence levels (top), driven by reduced errors for negative valence under ketamine. A similar drug x emotion interaction was trending for arousal (bottom; p = .051). At medium arousal levels, ratings were shifted towards higher arousal following ketamine. **C.** Integration kernels for valence (top) and arousal (bottom) by drug and volatility. Lines show mean kernel weights (dashed, placebo; solid, ketamine); shaded bands ± within-subject s.e.m. (Cousineau-Morey). In valence, following ketamine, kernels were broader in stable blocks, with relatively more weight on the second most recent face, and steeper in volatile blocks, with more weight on the most recent face. In placebo sessions, kernel shape was similar across volatility conditions. Arousal integration kernels did not differ between drug conditions. **D.** Integration kernel steepness (IKS), by drug and volatility. In valence, the effect of volatility differed between the two drug conditions (drug x volatility interaction effect, p = .016): IKS was higher in volatile than in stable blocks following ketamine (Holm p = .007). Arousal IKS was not affected following ketamine. Black dots and error bars denote group level means ± s.e.m. with Cousineau-Morey correction, whereas coloured dots denote participant level data. ** p < .01, * p< .05, † p < .10.

Ketamine interacted with several measures of task performance but did not produce global changes in task engagement. There were no overall effects of ketamine on response frequency, change-point detection accuracy, absolute rating errors, or reaction times in either valence or arousal (all p > .1; **Fig S5**). In arousal blocks, ketamine interacted with volatility, producing small reductions, specifically in volatile blocks, in both rating changes (drug x volatility interaction: b = -0.078, 95% CI [-0.154, -0.001], χ²(1) = 3.960, p = .047; follow-up volatile ketamine-placebo Δ = -2.16, Holm p = .029; **Fig S5B**) and change-point accuracy (drug x volatility interaction: b = -0.299, 95% CI [-0.515, -0.082], χ²(1) = 7.29, p = .007; follow-up volatile ketamine-placebo Δ = -0.035, Holm p = .045; **Fig S5C**). Ketamine also modulated rating errors across negative and positive valence levels (drug x emotion: b = 0.118, 95% CI [0.032, 0.204], F(1,636) = 7.216, p = .007), driven by reduced errors for negative valence under ketamine (Δ = -0.071, Holm p = .046). In arousal blocks there was a trending drug x emotion interaction (b = -0.123, 95% CI [-0.246, 0.001], F(1, 636) = 3.82, p = .051). Additionally, ketamine increased positive signed error at medium arousal levels (b = 0.115, 95% CI [0.015, 0.216], F(1,28) = 5.561, p = .026; **Fig 5B**), suggesting overestimation of medium arousal following ketamine. Thus, ketamine produced small, context-specific changes in behavioural performance and emotional biases rather than a general disruption of task performance.

To test whether ketamine altered the temporal weighting of emotional evidence we used the regression-based integration kernels. In valence, ketamine selectively modulated kernel shape as a function of environmental volatility (drug x volatility interaction: b = 0.077, 95% CI [0.015, 0.139], F(1, 84) = 6.03, p = .016; **Fig 5C-D**), with no main effect of drug (b = 0.004, 95% CI [-0.027, 0.035], F(1, 84) = 0.06, p = .806). Following ketamine, kernels were steeper in volatile than stable blocks (ketamine volatile-stable: b = 0.067, 95% CI [0.016, 0.117], t(84) = 3.01, Holm p = .007), whereas in placebo IKS did not differ between volatility conditions (placebo volatile-stable: b = -0.010, 95% CI [-0.061, 0.040], t(84) = -0.463, Holm p = .644). Neither drug comparison within a volatility condition was significant (stable ketamine - placebo: b = -0.035, 95% CI [-0.085, 0.016], t(84) = -1.562, Holm p =.122, volatile ketamine - placebo: b = 0.042, 95% CI [-0.008, 0.093], t(84) = 1.910, Holm p = .119). In arousal ketamine did not affect IKS (main effect of drug: b = 0.006, 95% CI [-0.018, 0.030], F(1, 84) = 0.25, p = .622; drug x volatility interaction: b = 0.001, 95% CI [-0.047, 0.050], F(1, 84) = 0.00, p = .960). The preregistered time-binned kernel did not show any drug or volatility effects on IKS, including the valence drug x volatility interaction (p = .223; **Fig S4C-D, Supplementary Table S3b**). Preregistered paired comparisons of stable block weights in the second most recent face did not reach significance in either rating dimension (beta weight in face -2, valence ketamine-placebo Δ = 0.019, 95% CI [-0.014, 0.051], t(28) = 1.18, p = .249; arousal, ketamine-placebo Δ = -0.003, 95% CI [−0.027, 0.020], t(28) = -0.30, p = .770).

Based on prior work linking NMDA receptor hypofunction to altered evidence accumulation and premature commitment to decisions under uncertainty (Cavanagh et al., 2020; Salvador et al., 2022), we preregistered the hypotheses that ketamine would increase recency weighting and reduce differences between stable and volatile environments. Neither preregistered prediction was supported. However, our findings diverge from these predictions in an interesting way. Rather than inducing a uniform shift towards recent evidence, ketamine changed how evidence integration depended on the environmental statistics: when rating valence, kernels were broader in stable blocks and steeper in volatile blocks following ketamine. This difference was not detected in the placebo condition. No drug effect on evidence integration was detected in arousal. This pattern suggests that ketamine alters how evidence is temporally integrated in a volatility context-specific manner, rather than globally increasing reliance on recent information.

## Discussion

Everyday social interactions require continuous integration of affective information in dynamic environments, without explicit cues indicating when to update beliefs or make decisions. Yet most experimental paradigms in emotion research have presented emotional stimuli as isolated, static events. Here, we introduce a novel task that captures emotional processing as a continuous process of evidence integration (Glaze et al., 2015; Huk et al., 2018; Hyafil et al., 2023) by requiring participants to integrate noisy affective cues over time to infer underlying emotional states. Using this approach, we showed that participants tracked dynamically changing emotional signals and were sensitive to the statistical structure of the environment. Tracking also showed systematic emotion-dependent biases: negative valence and high arousal were tracked more closely, and responses to changes towards them were faster.

Behaviour was sensitive to environmental volatility, but less clearly than predicted. In volatile blocks, participants changed their ratings more often and, when rating valence, responded faster than in stable blocks. The integration kernels showed steeper weighting of recent evidence in volatile blocks when rating valence in the online cohort, with a trend-wise effect in the same direction in the in-person cohort, but no volatility effect for arousal in either cohort. These patterns of behaviour are broadly consistent with principles established in perceptual decision-making, where evidence accumulation is flexibly adjusted according to environmental volatility (Gold & Shadlen, 2007; Ossmy et al., 2013; Ruesseler et al., 2023). Our results therefore extend computational accounts of perceptual inference to the affective domain, suggesting that similar principles may apply to the integration of emotional evidence.

We examined whether the adaptive adjustment of temporal evidence integration to environmental statistics is altered following ketamine administration. NMDA receptors play a central role in the cognitive inference processes required for evidence accumulation (Cavanagh et al., 2020; Salvador et al., 2022). During acute ketamine administration, healthy volunteers showed increased inference errors and premature commitment to decisions, despite largely preserved sensory processing (Salvador et al., 2022). In contrast, our findings in the post-acute phase point towards a more selective modulation of emotional evidence integration, rather than a global disruption of inference. Contrary to our preregistered prediction, ketamine did not increase the overall steepness of evidence integration. Instead, it changed the temporal weighting of emotional evidence in a context-dependent manner. In valence blocks, recent evidence was weighted more strongly in volatile than in stable blocks following ketamine, but not placebo. This pattern suggests that post-acute effects of ketamine may not simply disrupt inference but change how evidence is integrated over time depending on environmental structure. Within a predictive coding framework, reduced precision of prior beliefs could provide a common mechanism linking these effects to ketamine’s psychotomimetic and antidepressant actions, by weakening the influence of overly precise priors about the structure of the environment and allowing incoming evidence to exert greater impact on updating.

Ketamine is a rapid-acting antidepressant (Berman et al., 2000; Krystal et al., 2024; Sanacora et al., 2008), with clinical effects emerging within hours of administration in treatment-resistant depression (Zarate et al., 2006). Given that emotional processing is often perturbed in psychiatric disorders, most notably through negative biases in depression, several studies have investigated ketamine’s effects on emotion processing (Abel et al., 2003; Scheidegger, Henning, Walter, Boeker, et al., 2016; Scheidegger, Henning, Walter, Lehmann, et al., 2016; Schmidt et al., 2013). In healthy volunteers, ketamine reduced amygdala and limbic responses to fearful stimuli, and increased activity in the visual cortex in response to neutral stimuli (Abel et al., 2003). S-ketamine also reduced the discrimination of fearful and happy relative to neutral emotions, and reduced N170 amplitudes in response to emotional faces, while early sensory responses (P100) were preserved (Schmidt et al., 2013). This suggests that ketamine primarily affects the structural encoding of emotional faces rather than the earliest extraction of emotion-related visual information. Importantly, these studies have typically focused on acute effects during or shortly after infusion, although changes in affective processing have also been observed at later post-acute timepoints (Meiering et al., 2024; Pulcu et al., 2026; Reed et al., 2019). We tested participants approximately 1.5 hours after infusion, during the early post-acute period when antidepressant and plasticity-related effects may be emerging (Duman et al., 2016; Krystal et al., 2024; Zarate et al., 2006). Our findings suggest that, at this stage, ketamine’s effects on emotion processing may extend beyond responses to individual stimuli to how emotional evidence is integrated over time.

In our task, ∼1.5 hours following the infusion, ketamine did not produce a general disruption of task performance but instead had small and context-specific effects. In arousal blocks, ketamine reduced rating changes and change-point accuracy specifically in volatile environments. Ketamine also modulated emotion-specific rating biases, with reduced errors for negative valence and higher ratings of medium arousal. The arousal performance effects may reflect the greater perceptual ambiguity of arousal (Calvo & Nummenmaa, 2016; Mattek et al., 2017), which may become particularly consequential in volatile environments that require more frequent updating. This increased uncertainty may interact with ketamine’s effects on inference processes, leading to less accurate belief updating under rapidly changing conditions. This interpretation is consistent with prior work showing that ketamine impairs inference under uncertainty while leaving basic sensory processing relatively intact (Salvador et al., 2022). Beyond these effects on task performance and rating biases, ketamine selectively modulated the temporal integration of emotional evidence, suggesting changes in how evidence is accumulated over time that are not fully reflected in final emotion ratings alone. This may reflect a different aspect of emotion processing from that captured in discrete emotion recognition paradigms, as our task places greater emphasis on integrating information over time rather than evaluating isolated stimuli. Given that participants continuously integrate evidence across multiple faces rather than making discrete choices after each stimulus, ketamine-related changes in emotional processing may be more readily detectable in face-wise neural responses than in overt behaviour. This should be addressed in future work focusing on neural markers of emotional evidence integration.

The present study has several limitations. First, our study was conducted in healthy volunteers using a three-stage infusion protocol (a 10-min bolus followed by stabilisation and maintenance phases; approximately 0.71mg/kg over 90 minutes; see Methods) designed to maintain a stable plasma concentration. This differs from the 0.5mg/kg, 40-min infusion commonly used in clinical studies of ketamine’s antidepressant effects (Berman et al., 2000; Zarate et al., 2006), although higher doses and extended infusions have also been reported to have clinical benefit (Sanacora et al., 2017). The differences in protocol and population may limit the generalisability of the findings to clinical populations. Previous work has shown that ketamine can produce qualitatively different effects on emotional processing in healthy individuals compared to patients with depression, including opposite patterns of change in behavioural and neural responses (Nugent et al., 2019; Reed et al., 2019). Second, the task was completed 1.5 hours after the end of the infusion, coinciding with the onset of the putative neuroplasticity window (Duman et al., 2016; Krystal et al., 2024) and the earliest antidepressant effects, although the peak of ketamine’s antidepressant effects is thought to occur around 24 hours later (Zarate et al., 2006). This timing makes it difficult to attribute the observed behavioural effects to a specific mechanism, as residual NMDA receptor antagonism may overlap with downstream glutamatergic, plasticity-related, and circuit-level processes initiated by ketamine (Dai et al., 2025; Jelen et al., 2026; Krystal et al., 2024; Ma et al., 2023). It is therefore possible that ketamine’s effects on emotional biases and evidence integration differ across timepoints as the relative contribution of these mechanisms changes (Reif-Leonhard et al., 2025). Future studies comparing clinical and non-clinical samples, and examining multiple post-infusion timepoints, will be important for clarifying how the present findings relate to ketamine’s antidepressant effects. Third, while our task captures continuous emotional evidence integration, we did not include a directly comparable perceptual evidence accumulation task in the post-acute phase of ketamine administration. This limits our ability to determine whether the ketamine effect on emotional evidence integration reported here is specific to emotional stimuli or reflects a broader effect on evidence accumulation. However, because valence and arousal were rated on the same stimulus sequences within the same task structure, a general effect on evidence accumulation would be expected to affect both dimensions similarly. That ketamine modulated evidence integration when rating valence but not arousal therefore raises the possibility that its effects may not be entirely nonspecific, even within emotional processing, although differences in the difficulty of arousal judgements may also contribute. Fourth, as a novel continuous paradigm, our task required analytic choices that trial-based tasks avoid, such as response window definitions, how consecutive button presses are merged, and how integration kernels are specified. We based these choices on task structure and response distributions, but alternative choices are possible. In particular, because stable and volatile change-point intervals did not overlap, part of the RT difference between environments may reflect the shorter observation window in volatile blocks (**Fig S3E**). We also note that the use of discrete rating scales may have introduced ceiling effects and limited our sensitivity to subtle changes in behaviour, particularly at extreme levels of valence and arousal (**Fig S2D**). Finally, as is common in studies involving drugs with noticeable subjective effects, maintaining blinding was challenging. Despite the double-blind design, all participants who completed a retrospective debrief correctly identified the ketamine session (see Supplementary Results). Although the task was performed after the acute subjective effects had largely subsided, we cannot exclude the possibility that awareness of treatment allocation or associated expectations influenced task behaviour. However, visit order was counterbalanced, and the drug effects were unchanged when the preregistered covariates (visit order and gender) were included (**Supplementary Table S9**).

Together, these findings highlight the importance of considering emotional processing as a dynamic and context-dependent process that unfolds over time. By combining a continuous behavioural paradigm with pharmacological manipulation, the present study provides a novel framework for investigating how affective information is integrated in changing environments. Our results suggest that, in the post-acute phase, ketamine does not globally disrupt affective tracking, but has small, context- and emotion-specific effects, including a change in how the temporal integration of emotional evidence depends on environment volatility. Speculatively, more flexible integration of new emotional evidence could be one route through which ketamine exerts its rapid antidepressant effect and reduces the impact of maladaptive negative biases in depression, a hypothesis that remains to be tested in clinical samples. More broadly, continuous paradigms provide a tool for understanding whether emotional disturbances in depression and other mood disorders reflect not only biased evaluation of individual emotional signals but also altered integration of emotional evidence over time.

## Methods

### Participants

Forty-eight healthy volunteers attended the screening visit for the in-person study. Of these, three did not meet the eligibility criteria, three withdrew before the first infusion visit, and in two intravenous cannulation was not possible. Of the 40 participants who received an infusion, the first three were used to pilot the newly developed tasks and the infusion protocol and were not included in the analyses. Seven participants were excluded due to ketamine-related adverse effects. In two, the infusion was terminated during the first infusion visit (one because of severe nausea, one because of a hypertensive episode), and they did not complete the study. The remaining five completed both visits, but moderate to severe nausea and vomiting required the infusion to be temporarily interrupted to administer ondansetron, and this interruption resulted in non-standard ketamine exposure. These side effects also impaired their performance on the perceptual decision-making task administered during the infusion. In all cases, symptoms resolved shortly after the infusion was stopped. One additional participant was excluded due to failing preregistered behavioural quality checks on the continuous emotion rating task (task disengagement characterised by too few rating changes during task blocks). The final sample therefore comprised 29 participants (15 female, aged 18-40 years, M = 24.79, SD = 5.46). Participants were paid £250 for completing the study and a bonus of up to £50 depending on task performance. The study was approved by the University of Oxford Central University Research Ethics Committee (CUREC; Ref: R77437/RE001), and all participants provided written informed consent. Hypotheses and analyses were preregistered on AsPredicted following completion of data collection but prior to unblinding of drug conditions (https://aspredicted.org/bp4w-9t7p.pdf).

An independent online sample of self-reported healthy volunteers was recruited via Prolific for a single online session of the same task. Participants were paid £7 and up to £3 bonus depending on task performance. Of the 152 participants who completed the task, 60 were excluded for failing behavioural quality checks (the same criteria as preregistered for the in-person study). The final sample comprised 92 participants (32 female, aged 20-40 years, M = 31.46, SD = 5.45). All participants provided informed consent online. The online study was approved by the Central University Research Ethics Committee (CUREC; Ref: R73912/RE001).

### Continuous emotion rating task

We developed a novel continuous emotion rating task to examine how affective evidence is integrated over time and how integration is adjusted depending on environmental statistics. The task was programmed in PsychoPy v2022.2.5 (Peirce et al., 2019); the online version ran in PsychoJS and was hosted on Pavlovia (https://pavlovia.org/). In this task, participants viewed a rapid stream of facial stimuli and continuously rated the average level of either emotional valence or arousal. The face stimuli were selected from the KDEF-dyn database (Calvo et al., 2018), a dynamic version of the Karolinska Directed Emotional Faces (KDEF) database (Lundqvist et al., 1998), in which neutral faces are morphed into expressive faces (e.g., happy, angry, fearful, disgusted) to produce a continuous range of emotional intensities. Valence and arousal values for each morph image (31 morph levels per expression) were obtained by linear interpolation between the normative ratings of the neutral and full-expression images provided with the database (range: valence 1.0-7.8, arousal 1.4-7.3). These values were used to generate moment-to-moment face stimulus sequences, while the mean level was constrained to a smaller set of values (2-6 in steps of one for valence and 2-6 in steps of two for arousal). Since it is difficult to distinguish fine levels of arousal, participants reported their ratings on a five-point scale for valence (very negative to very positive) and a three-point scale for arousal (low to high). Facial stimuli were presented for an average of 0.36 seconds (range: 0.02-0.7 seconds). Due to an error in sequence generation, about 5% of faces were shown for less than 0.2s and were excluded from the integration kernel analysis. The cover story for this task was that participants were rating the average level of emotions expressed by an audience watching two different movies.

To examine how emotional evidence is integrated over time in different environments, and how this might be shaped by expectations or priors, we introduced several key manipulations. First, faces were presented with levels of valence or arousal that were noisy samples drawn from a normal distribution centred around a true underlying mean. The underlying mean shifted unpredictably at certain intervals, generating change-points of varying magnitude. For valence, changes could be small (1-step, corresponding to one scale step), medium (2-step), or large (3-step); for arousal, they could be small (2-step, corresponding to one scale step, e.g. low to medium) or large (4-step, corresponding to two scale steps, e.g. low to high). Participants’ task was to track this latent mean as accurately as possible by updating their ratings.

Second, we manipulated the volatility of the environment across two conditions, stable and volatile. Each condition was cued by a distinct frame shape (square or round frames for Movies A and B in visit 1, triangular or hexagonal frames for Movies C and D in visit 2). Stable blocks were characterized by longer intervals between change-points (7.27-11.10 seconds, median = 8.4s), whereas volatile blocks had shorter intervals (3.10-6.95 seconds, median = 4.3s). The variability of the individual face values around the underlying mean was the same in both conditions (SD = 0.8). During the three-day online training period (see below), participants learned through experience that the two contexts (“movies”) differed in the frequency with which the underlying emotional state changed. They were not explicitly informed which context was stable or volatile. Instead, the distinct frame shapes identified the two contexts, allowing participants to learn an association between each contextual cue and its volatility. During subsequent task performance, participants could therefore use these learned contextual expectations while also continuing to infer volatility from the incoming evidence.

Training consisted of a structured progression to familiarise participants with the temporal dynamics of the task and to induce priors about environmental volatility. Participants first watched a video showing the slider automatically tracking the true underlying mean in Movies A and B (Movies C and D before the second visit; training was otherwise identical in both weeks), to give them a sense of the frequency of changes. They then completed two training blocks (one stable, one volatile) with immediate feedback on their ratings, in which the frame surrounding each face turned green when the participant’s rating matched the true mean and red otherwise. This colour feedback was removed in subsequent training blocks. Across the three days of online training, participants completed four valence blocks (two stable, two volatile; two with colour support, two without) on day 1, four arousal blocks (same structure) on day 2, and four blocks without colour support (mixed valence/arousal) on day 3. During the infusion sessions, all blocks were performed without feedback (but with shape cues to indicate the movie). At the end of each block, participants received summary feedback comparing their performance (mean rating error) with that of an independent online pilot sample (n=20), which was collected prior to the in-person study to provide a performance baseline, and which was otherwise not analysed here.

The main task was conducted 1.5 hours after the end of the infusion (M = 93 minutes, range of task start times 71-122 minutes) to examine ketamine’s delayed effects on emotional integration kernels, coinciding with the expected onset of its antidepressant action reported in clinical populations. Participants completed 12 blocks, 2.5 minutes each, evenly divided between valence and arousal ratings, and volatility conditions, the order counterbalanced across participants (4 orders). The same 6 stimulus sequences (3 stable and 3 volatile) were used for all participants. Each sequence was rated once for valence and once for arousal in each visit. Twice per block (midway through and at the end), the face stream was paused and participants rated their confidence in their current rating, followed by their own current valence and arousal on the same scales as the task (2s response limit for each question).

### Study design and timeline

The study used a randomised, double-blind, placebo-controlled crossover design. Participants completed two infusion visits one week apart, with the one-week interval providing a washout period to minimise potential carryover effects between drug conditions. Participants received intravenous racemic ketamine during one visit and saline placebo during the other. Treatment order was randomised and stratified by gender to balance ketamine-placebo and placebo-ketamine sequences among male and female participants. The randomisation list was generated using block randomisation with an online tool (https://www.sealedenvelope.com/simple-randomiser/lists), by a researcher not otherwise involved in the study. Participants were allocated to a treatment order after screening and before their first infusion visit. Participants and researchers conducting the behavioural assessments were blinded to treatment condition, while the physician and research nurse responsible for drug administration and clinical monitoring were unblinded. Participants’ perception of treatment allocation was assessed retrospectively in an end-of-study debrief questionnaire (for details see Supplementary Results).

Participants first completed a screening visit at the NIHR Clinical Research Facility: Oxford Health, Warneford Hospital, Oxford. Screening included medical history, psychiatric assessment using the Structured Clinical Interview for DSM-5 (SCID-5), physical examination, electrocardiography, blood pressure measurements, blood tests (full blood count, glucose, liver function, urea and creatinine, thyroid-stimulating hormone) and urine tests (drug screen and, where applicable, pregnancy test). Participants were eligible if they were aged 18-40 years, had normal or corrected-to-normal vision, had no current or past psychiatric or neurological disorder, had not previously used ketamine or phencyclidine, and were not pregnant or breastfeeding. Eligible participants were subsequently scheduled for the two infusion visits.

Three days before each infusion visit, participants began online training on the continuous emotion rating task to familiarise them with the task and establish prior expectations about the environmental conditions (see Continuous emotion rating task). Participants also trained on two additional tasks that are not reported here.

At the beginning of each infusion visit, continued eligibility was confirmed by clinical staff, including review of concomitant medications and urine toxicology and, where applicable, pregnancy testing. Participants were required to fast on the day of the infusion before drug administration and were given lunch after the end of the infusion, before the continuous emotion rating task. Participants completed behavioural tasks and questionnaires before, during, and after drug administration. Psychotomimetic, dissociative, and affective effects were assessed using the Brief Psychiatric Rating Scale (BPRS), simplified 6-item Clinician-Administered Dissociative States Scale (CADSS-6), and Positive and Negative Affect Schedule (PANAS). BPRS and CADSS were administered before, during, and after infusion, while PANAS was administered before and after infusion.

The continuous emotion rating task was performed approximately 1.5 hours after the end of the infusion. EEG was recorded during task performance but is not reported in the present study. Participants remained under clinical observation following the infusion and were discharged by a physician once considered clinically fit to leave. Following discharge, participants had access to a 24-h contact number for medically qualified on-call staff.

### Infusion protocol

Racemic ketamine or saline solution was administered intravenously using a previously described three-stage protocol (Salvador et al., 2022): a 10-minute bolus (0.023 mg/kg/min), a 20-minute stabilisation phase (0.0096 mg/kg/min) and a 60-minute third lower dose phase (0.0048 mg/kg/min), corresponding to a total dose of approximately 0.71 mg/kg. The three-stage protocol was designed to achieve a stable ketamine plasma concentration of approximately 150 ng/ml during a perceptual decision-making task performed during the infusion, which is not reported in the present study. Blood pressure, heart rate, and peripheral oxygen saturation were monitored at 10-min intervals throughout the infusion. Adverse effects were managed clinically as required, including temporary interruption or termination of the infusion and administration of ondansetron for nausea. Seven participants experienced adverse effects requiring interruption or termination of ketamine administration and were excluded from the final sample as described above.

### Behavioural analysis

To evaluate performance on the continuous emotion rating task, we quantified three complementary measures: a continuous framewise measure of tracking accuracy (rating error), and change-point (CP)-based measures capturing detection accuracy and response times (RTs) for correctly detected change-points (correct RT). Emotion level was defined based on the true underlying mean, with valence grouped into negative (2-3), neutral (4), and positive (5-6), and arousal into low (2), medium (4), and high (6). Framewise signed rating error was computed as the difference between the participant’s rating and the true underlying mean (participant rating - true mean). To allow comparison of error direction across the extreme levels (negative/positive valence and low/high arousal), the sign of the rating error was flipped for positive valence and high arousal, such that larger values indicated ratings that were less extreme than the true mean (i.e. closer to the centre of the scale).

CPs were defined as frames where the true underlying mean changed its value. For each CP we defined a valid response detection window that started after a short buffer (1 s for the in-person cohort; 0.5 s online) to avoid counting responses still attributable to the preceding CP, then extended until the earliest of: the next CP plus the same buffer, 7s after CP onset, the mid-block confidence check or the end of the block. The 7s limit prevented excessively long detection windows in stable blocks and lies just below the shortest stable CP interval (7.3s) and above the longest volatile CP interval (6.95s). We excluded CPs whose detection window was still open at the mid-block confidence check, as well as the CPs at the end of the block with detection windows shorter than 1s (i.e. CPs shorter than 2s in person and 1.5s online, at the end of the block). Buffer durations were supported by the inspection of the RT histogram and the detection time course, which showed a sharp increase at approximately 1 second after a change-point for the in-person sample (**Fig 2A**) and earlier at around 0.5s for the online cohort (**Fig S1A**). Few change-points were detected within the first second, with most of these early responses in the same direction as the previous change-point, indicating that they partly reflected delayed responses to the previous change. A CP was scored as correctly detected if the participant made a rating change in the direction of the true mean jump within the detection window. Correct RT was measured as the latency from the CP frame to the first correct participant rating change and only correct CPs contributed to RT analyses. For comparisons between emotion levels, analyses were restricted to CP magnitudes 1-2 (excluding large CP magnitudes), because larger CP magnitudes cannot result in transitions towards intermediate emotion levels (neutral valence and medium arousal), leading to imbalances across conditions. Within each rating dimension, emotion levels were compared within change-point direction: for valence, negative versus neutral following downward changes and neutral versus positive following upward changes; for arousal, low versus medium following downward changes and medium versus high following upward changes (**Fig 3C**).

To illustrate how quickly participants responded to changes in the true underlying mean, we estimated the cumulative proportion of CPs detected as a function of time since CP onset. A CP was considered detected at the first rating change in the direction of the underlying mean shift. Because CP intervals differed between volatility conditions (stable 7.3-11.1s, volatile 3.1-6.95s), many CPs, particularly in volatile blocks, ended before a slow response could occur. We therefore used the Kaplan-Meier estimator (E. L. Kaplan & Meier, 1958), as in event-history analyses of response times (Jahn-Eimermacher et al., 2011; Panis & Ramsey, 2026). CPs which ended before a response was made were treated as censored instead of counted as a miss.

We compared participant task performance (rating error and change-point accuracy) with a null model to assess whether responses reflected meaningful tracking of the underlying stimulus rather than random behaviour. Null datasets were generated by shuffling contiguous rating runs within each block for each participant, while preserving the original run durations and the distribution of time spent at each rating level. This preserved the temporal structure of participants’ responses while disrupting their alignment with the underlying emotional signal. For each participant block, the shuffling procedure was repeated 1000 times, and the same performance metrics were computed for each shuffled dataset (**Fig 2BC**).

Mixed-effects models were used for statistical inference and were fitted in R using lme4 (Bates et al., 2015), lmerTest (Kuznetsova et al., 2017), pbkrtest (Halekoh & Højsgaard, 2014), and emmeans (Lenth et al., 2026). Change-point (CP) detection accuracy was analysed at the CP level using binomial GLMMs, while log-transformed reaction times (RT) and rating errors (RE) were analysed using linear mixed-effects models (LMMs). Rating-error analyses used subject x drug x dimension x block x emotion-level means. IKS analyses used condition-level estimates, averaging the three repeated same-condition blocks within each subject x drug x dimension x volatility cell. Categorical predictors were coded using sum-to-zero contrasts. Binary predictors (drug: placebo/ketamine; volatility: stable/volatile; dimension: valence/arousal; CP direction: downwards/upwards) used -0.5/+0.5 coding, such that coefficients reflect the second level minus the first. CP magnitude was centred around the midpoint of the possible magnitude values within each dimension (valence CP magnitude - 2; arousal CP magnitude - 3). Random-effects structures were selected using a parsimonious mixed-model strategy following Bates et al. (Bates et al., 2018). We started from the maximal subject-level random-effects structure justified by the design and simplified singular or overparameterized models by first removing random-effect correlations and then unsupported random slopes, until the fit was non-singular. For Gaussian LMMs, fixed-effect inference used Type-III Kenward-Roger F tests. For binomial GLMMs, inference used Type-III likelihood-ratio tests, with Wald z tests for intercepts. Coefficients are reported on the log-odds scale, and follow up contrasts for accuracy are reported as differences in probability. Significant effects were followed up with pairwise comparisons using estimated marginal means (emmeans) with Holm correction, Kenward-Roger degrees of freedom and Bonferroni-adjusted confidence intervals. For follow-up analyses separating change-point direction from reached emotion level, models were fitted separately within each direction and dimension. Because reached emotion level and CP magnitude were not fully crossed, these tests were restricted to the common CP-magnitude support for the compared emotion levels, with CP magnitude included as a covariate where applicable. All tests were two-sided. Error bars show within-subject standard error of the mean computed by normalising each participant’s data to the grand mean (Cousineau, 2005) and applying Morey’s correction (Morey, 2008).

We quantified how evidence from recently seen faces influenced rating with regression-based integration kernels (Brunton et al., 2013; Keung et al., 2019; Wyart et al., 2012). For each block we first computed framewise prediction errors (PEs) as the difference between the moment-to-moment face stimulus valence or arousal level and the participant’s current rating. Each block was split at the mid-block confidence check so that no regressor spanned the pause. Rating changes were grouped into movement episodes to account for large updates expressed through multiple rapid button presses. Consecutive rating changes were merged into one movement episode if each occurred within 40 frames (667ms at 60Hz) of the previous one. Each movement episode was anchored at its first step, and its outcome was the net number of signed steps (clipped at ±2). Episodes whose steps cancelled out (e.g. an up step immediately followed by a down step, net change 0) were discarded. Each movement episode was assigned to the face on the screen when it began. The regressors of a movement were the PEs of the five most recent faces preceding it. Evidence in the 200 ms (12 frames) before movement onset was excluded, because it is unlikely to influence a movement that is already being initiated. Faces with 200 ms (12 frames) or less of exposure outside this excluded window were skipped. This removed the very brief faces in the schedule (1-12 frames/17-200 ms, about 5% of face stimuli; see Continuous emotion rating task), as well as faces that were mostly covered by the excluded window. Every face during which no movement occurred was used as a no-movement observation (outcome 0), with the PEs of the five preceding faces as regressors. For each block, the signed outcome (-2 to +2) was regressed on the five face PEs and intercept by weighted least squares. Because no-movement observations greatly outnumbered movements (about 28 movements for 334 no-movement per block on average), observations were weighted so that both classes contributed equally (movement observations N / (2N_move_), no-movement observations N / (2N_no-move_), with N the number of observations in the block). The resulting regression coefficients (beta weights) from the most recent face to the fifth most recent form the integration kernel. Block weights were averaged over the three same condition blocks of each participant x drug x rating dimension x volatility. As preregistered (see preregistration https://aspredicted.org/bp4w-9t7p.pdf), kernel shape was summarised by the integration-kernel steepness (IKS), the difference between the weights of the most recent and second most recent evidence, and by the weights of the second most recent evidence (BW-2).

We also estimated the preregistered time-binned logistic regression-based integration kernels, which are reported as a secondary analysis (**Fig S4, Supplementary Table S3b**). During model development, after unblinding but before examining drug effects, we found that this design had two limitations. First, the bin width is a free parameter which strongly influenced the effect of volatility on kernel steepness. Second, adjacent bins were highly correlated, especially at smaller bin widths. In addition, a binary outcome (movement vs no-movement) discards the direction and size of rating changes, although change-points of different magnitudes required updates of one or several rating steps. We therefore used the face-wise linear kernel described above as the primary analysis instead of this preregistered kernel (for details see **Supplementary Deviations from preregistration**). In the preregistered time-binned logistic regression kernels (secondary analysis), we modelled how PEs preceding each face onset influenced the probability of updating the rating. PEs preceding each face onset were grouped into five consecutive 500 ms (30-frame) time bins extending backwards from stimulus onset. The dependent variable was whether a rating update occurred within 717 ms (43 frames, the longest face duration in the schedule) following the current face stimulus. Rating updates were coded as a binary outcome, with 1 indicating an update and 0 indicating no update. We used the same merging rule for consecutive movements as above, and the merged update took the net direction of its steps. Direction-aligned signed PEs were used, such that PEs associated with downward rating changes were sign-flipped, ensuring that positive values always reflected evidence favouring a rating update in the observed direction. Evidence integration kernels were estimated separately for each block using a class-weighted logistic regression (same weights as above) and IKS was computed in the same way but with the two most recent time-bin regressors instead of face regressors.

## Data availability

The behavioural results, debrief and mood questionnaire data will be made available upon publication.

## Code availability statement

Analysis code to reproduce all behavioural results will be made available upon publication.

## Acknowledgements

MCKF was funded by a Wellcome Trust Henry Dale Fellowship (223263/Z/21/Z), a UKRI-converted ERC Starting Grant (EP/X021815/1) and a Leverhulme Award in Psychology. LTH was funded by a Wellcome Trust Henry Dale Fellowship (208789/Z/17/Z). This study was supported through the National Institute for Health and Care Research (NIHR) Biomedical Research Centre (BRC). CZ, KAS and SA are supported by the National Institute for Health Research (NIHR) Oxford Health Clinical Research Facility (grant NIHR203975). We thank Rupert McShane for advice on the ketamine infusion protocol and for providing on-call clinical support during infusion visits. We thank Maja Friedeman for assistance with data collection, and Balaji Rajendran, Francisca Kaife-Nwokolo, Andrea Paggi, James Hong, Rosario Aronica and the wider team at the NIHR Oxford Health Clinical Research Facility (CRF) for their invaluable support in conducting the study. The views expressed are those of the authors and not necessarily those of the UK National Health Service, the NIHR, or the UK Department of Health.

For the purpose of open access, the author has applied a CC BY public copyright licence to any Author Accepted Manuscript version arising from this submission.

## Author contributions

MR, LW, LH, and MCKF conceived the study and designed the task. LH and MCKF secured funding for the study. MR and LW collected and analysed the data. PC, SC, CZ, and KAS advised on the clinical design, planning, and regulatory aspects of the ketamine study. CZ, SA and KAS supported drug administration and participant monitoring. JG and MCKF advised on data analyses. MR wrote the first draft of the manuscript, and all authors reviewed and edited the manuscript.

