## Supplemental Information for "Context-dependent effects of ketamine on emotional evidence integration"

### Supplementary Figures

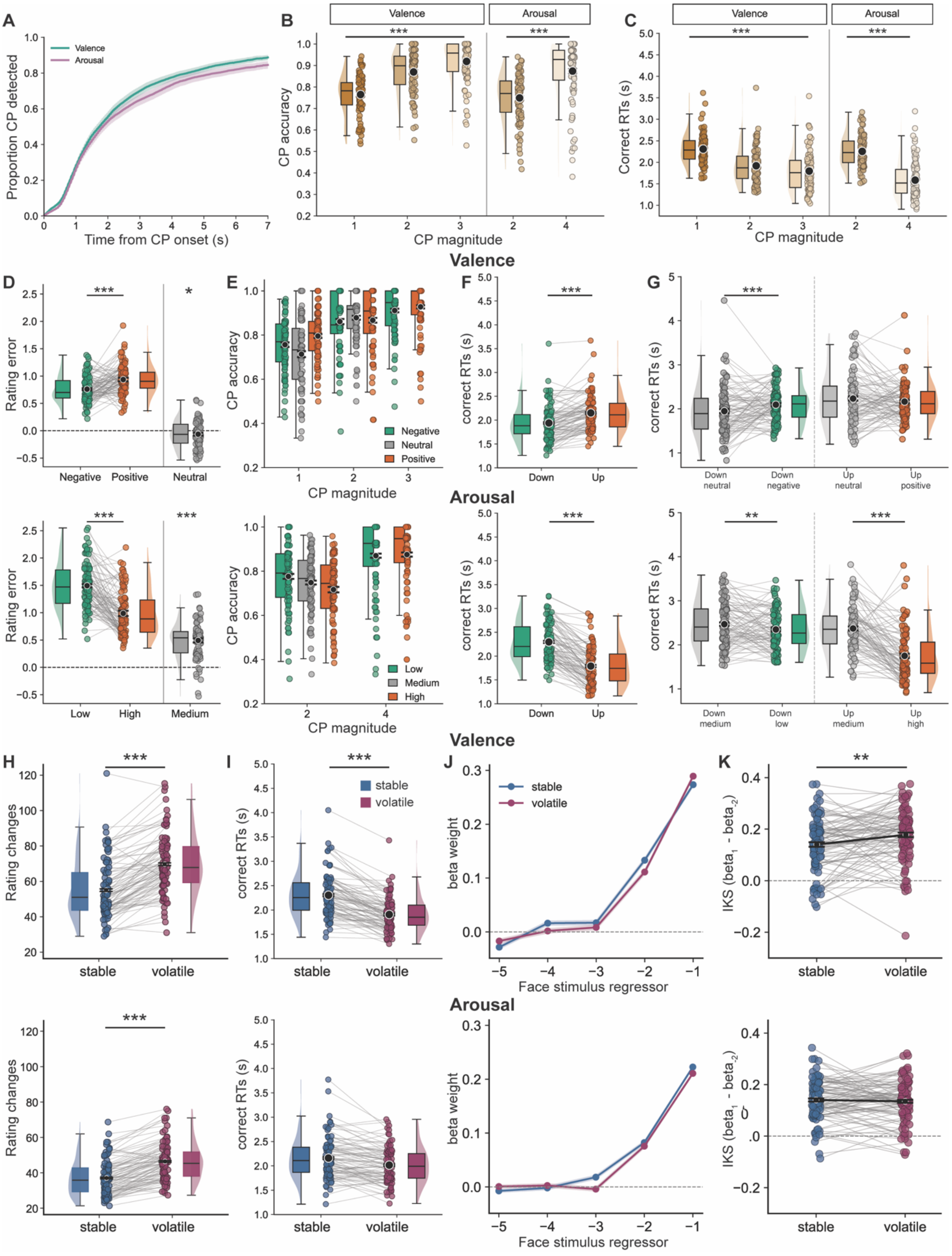


***Figure S1. Replication of main behavioural task effects in the online sample.*** Behavioural performance in the online sample (n=92) replicated the main task effects observed in the in-person sample (n=29). **A.** Cumulative proportion of detected change-points (CPs) as a function of time from CP onset, split by rating dimension (Kaplan-Meier estimate, mean ± 95% CI across participants).

Detection began to rise approximately 0.5s after CP onset, and half of all CPs were detected within approximately 1.8s. Valence showed slightly higher detection than arousal. **B.** CP detection accuracy increased with change-point magnitude for both valence (left) and arousal (right), indicating greater sensitivity to larger shifts in the underlying emotional true mean. **C.** Reaction times (RTs) for correctly detected CPs decreased with larger change-point magnitudes, showing faster responses to stronger changes in emotional evidence. **D.** Rating error (participant rating minus true underlying mean), across emotion levels. For comparability, rating error for positive valence and high arousal was sign-flipped. Tracking errors were lower for negative than positive valence (upper panel), and lower for high than low arousal (lower panel). Neutral valence was underestimated, whereas medium arousal was overestimated. **E.** CP detection accuracy split by the emotional level reached after the change and CP magnitude. In valence blocks, small changes towards positive valence were detected more accurately than changes towards negative or neutral valence, whereas at larger magnitudes accuracy was similar across emotion levels. In arousal blocks, small changes towards low arousal showed the highest accuracy, whereas larger changes towards both low and high were detected similarly well. Neutral valence and medium arousal cannot be reached by the largest CP magnitudes. **F.** RTs for correctly detected CPs split by CP direction. Participants responded faster to downward than upward changes in valence, and to upward than downward changes in arousal. **G.** RTs by CP direction and emotion level reached after the change, excluding the largest CP magnitudes which cannot reach intermediate levels. In valence blocks, responses after downwards changes were faster towards neutral than towards negative valence, whereas RTs did not differ between positive and neutral targets after upward changes. In arousal blocks, RTs differed more strongly by the emotion level reached, with slower responses towards medium than towards low or high arousal. **H.** Participants made more rating changes in volatile than stable blocks in both dimensions. **I.** Correct RTs split by volatility condition. Participants responded faster in volatile than stable blocks in valence, but not in arousal. **J.** Face-wise linear regression-based integration kernel beta weights of the prediction errors of the five faces preceding a change in rating, by volatility. Across both dimensions, recent prediction errors were weighted most strongly, indicating recency-weighted evidence integration. **K.** Integration-kernel steepness (IKS), defined as the difference between the beta weights of the two most recent face regressors ($beta_{-1}$ - $beta_{-2}$). Larger values indicate steeper, more recency-focused kernels. IKS was greater in volatile than stable valence blocks, consistent with shorter integration windows under greater volatility. Arousal blocks did not show an IKS difference between the two volatility conditions. Black dots and error bars denote group means ± within subject s.e.m. (Cousineau-Morey correction), coloured dots denote participant means. *** $p < .001$, ** $p < .01$, * $p < .05$.

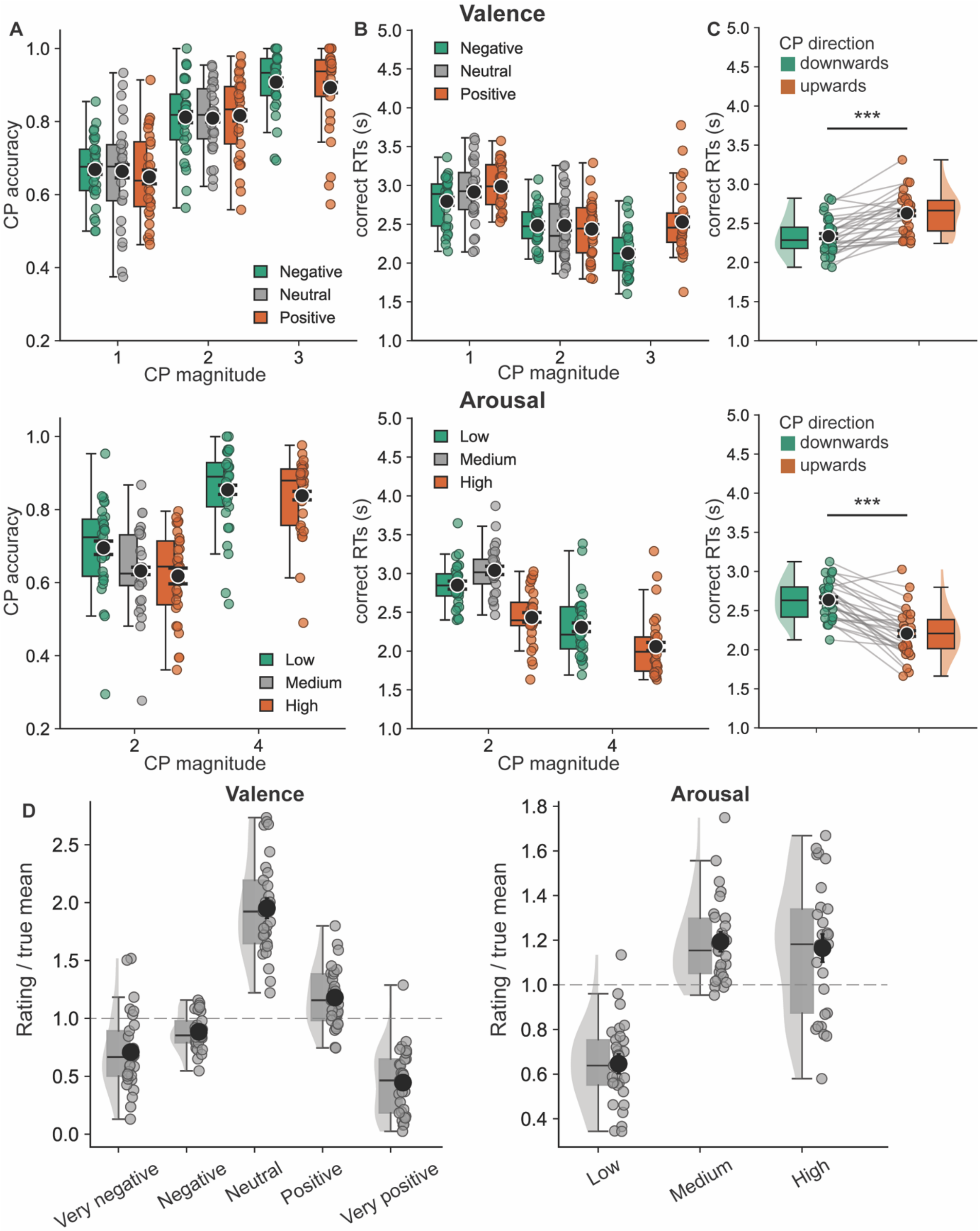


***Figure S2. Additional control analyses of emotional biases in the in-person sample.***
**A.** Change-point detection accuracy split by emotional level reached and change-point magnitude, shown separately for valence (upper panel) and arousal (lower panel) blocks. In valence blocks, accuracy was similar across emotion levels at each CP magnitude. In arousal blocks, in small jump sizes, changes towards low arousal showed the highest accuracy, followed by medium and high arousal, whereas large changes towards low and high arousal showed similar accuracy. **B.** RTs for correctly detected CPs split by emotional level reached and change-point magnitude. In valence blocks, responses were slowest for small changes towards positive levels and fastest for large changes towards negative levels. In arousal blocks, responses were faster for changes towards high arousal across change-point magnitudes. **C.** Control analysis of reaction times split by change-point direction, restricted to

downward changes reaching negative valence or low arousal and upward changes reaching positive valence or high arousal. Neutral valence and medium arousal were excluded because these levels could only be reached by smaller change-point magnitudes. This analysis showed the same direction as the full reaction-time analysis: participants responded faster to downward than upward changes in valence, and faster to upward than downward changes in arousal. **D.** Ratio of the number of frames participants spent at each rating level to the number of frames for which the true underlying mean occupied the corresponding level. Values above 1 indicate relative over-use of a rating level, whereas values below 1 indicate relative under-use. In valence blocks, participants disproportionately used the intermediate/neutral level, while under-using both extremes of the scale, consistent with compression of responses towards the centre. In arousal blocks, participants under-used the low-arousal level and over-used medium- and high-arousal ratings.

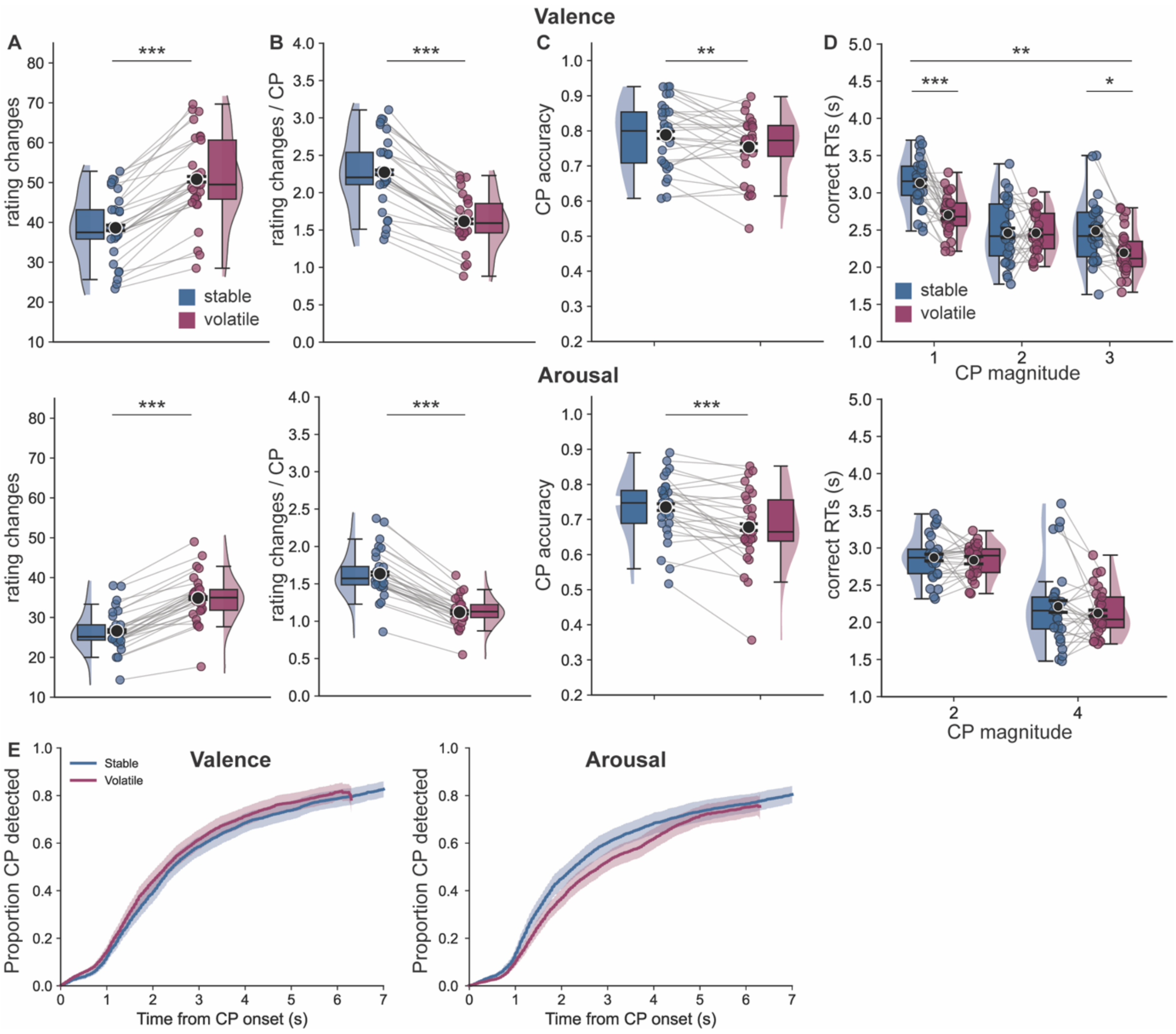


***Figure S3. Additional analyses of volatility effects and behavioural adjustments in the in-person sample***

**A.** Number of rating changes split by volatility condition, shown separately for valence (upper) and arousal (lower) blocks. Participants made more rating changes in volatile than stable blocks for both rating dimensions, consistent with the greater frequency of changes in the underlying emotional state in volatile environments. **B.** Ratio between the number of participant rating changes and the number of true change-points in each volatility condition. Ratios exceeded 1 in both stable and volatile blocks, indicating that participants made more rating changes than the minimum required to follow the true change-points. Participants made fewer ratings per change-point in volatile than stable blocks. **C.** Change-point detection accuracy split by volatility condition. Accuracy was higher in stable than volatile blocks for both valence and arousal, indicating that participants were more likely to correctly detect changes in the underlying emotional state when changes occurred less frequently, although the shorter response window in volatile blocks might also limit detection. **D.** Reaction times for correctly detected change-points split by volatility condition and change-point magnitude. When rating valence, participants responded faster in volatile than stable blocks for small (1-step) and large (3-step) changes, but not for medium (2-step) changes. No volatility effect was observed when rating arousal. **E.** Cumulative proportion of change-points detected as a function of time from change-point onset, split by volatility condition for valence (left) and arousal (right) blocks (Kaplan-Meier estimate, mean ± 95% CI across participants). Volatile curves end earlier because no volatile interval exceeded 7s. Black dots and error bars denote group means ± within subject s.e.m. (Cousineau-Morey correction), coloured dots denote participant means. *** p < .001, ** p < .01, * p < .05.

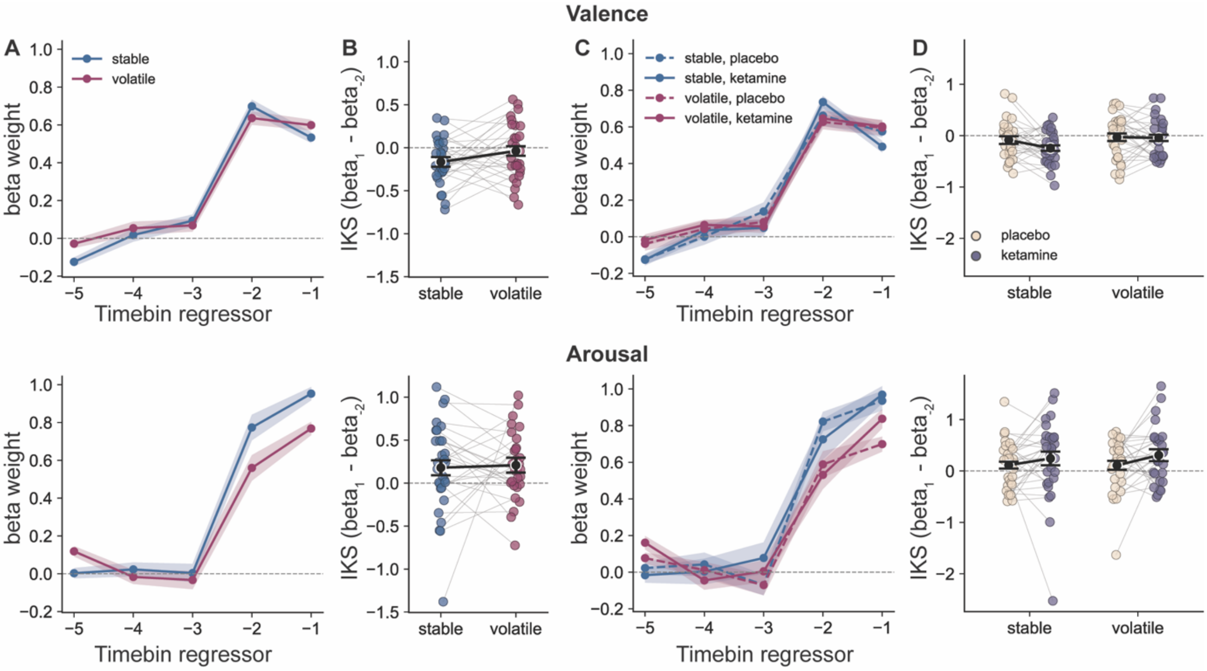


***Figure S4. Preregistered time-binned evidence-integration kernels (in-person cohort).*** Kernels were estimated with the preregistered time-binned logistic regression (five 30-frame / 500 ms bins of prediction errors preceding each face onset as regressors, predicting a rating update). Beta weights are on the log-odds scale and are plotted across the five preceding time bins. **A.** Kernels by volatility condition for valence (top) and arousal (bottom). As in the primary face-wise linear regression-based kernel analysis, recent prediction errors (the last two time bins, ~ 1s) contributed most strongly to subsequent rating updates. **B.** Integration-kernel steepness (IKS, difference between the beta weights of the most recent and second most recent time bin) by volatility condition. IKS did not differ significantly between stable and volatile blocks in either valence or arousal. **C.** Kernels by drug and volatility condition. **D.** IKS by drug and volatility condition. There was no significant effect of ketamine on IKS and no significant drug x volatility interaction for either valence or arousal. Lines and shaded bands show mean ± within-subject s.e.m, and coloured dots denote participant-level data.

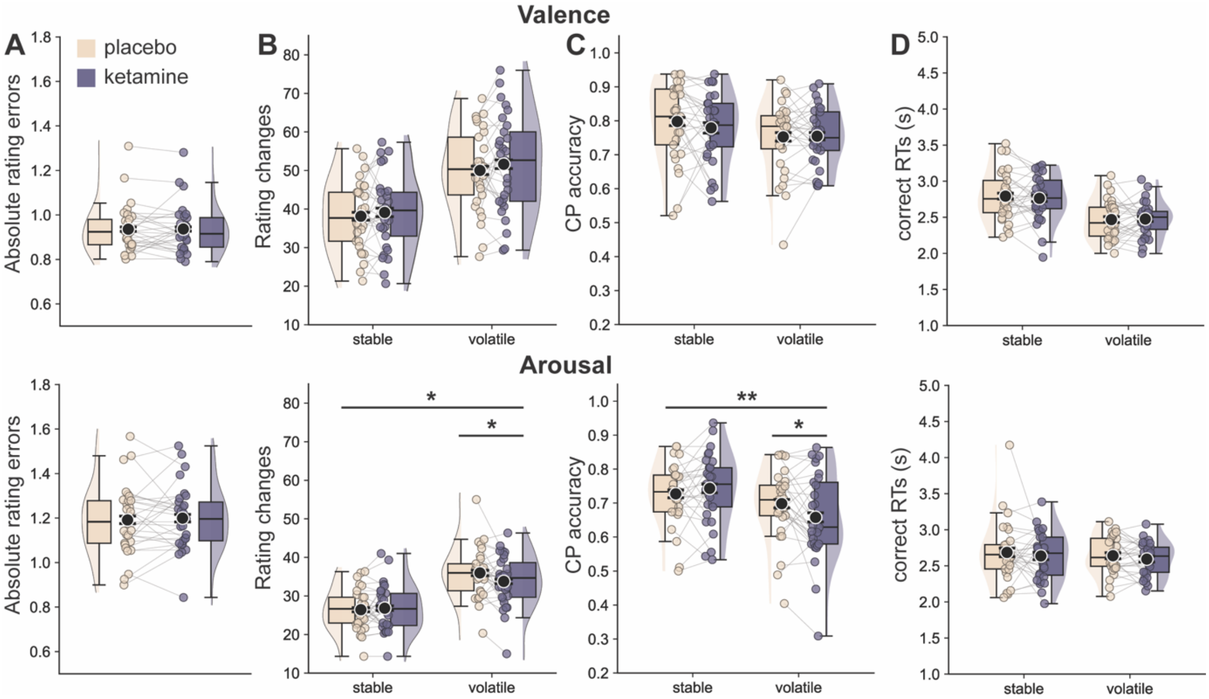


***Figure S5. Additional analyses of ketamine effects on behavioural adjustment in the in-person sample.*** **A.** Absolute rating errors under placebo and ketamine, shown separately for valence (upper row) and arousal (bottom row) blocks. Absolute rating errors did not differ between drug conditions. **B.** Number of rating changes split by drug and volatility condition. Ketamine did not affect response frequency in valence blocks, whereas in arousal blocks a small drug x volatility interaction reflected fewer rating changes following ketamine specifically in volatile blocks. **C.** Change-point detection accuracy split by drug and volatility condition. No ketamine-related differences were observed in valence blocks. In arousal blocks, a drug x volatility interaction reflected lower detection accuracy following ketamine in volatile but not stable blocks. **D.** Reaction times for correctly detected change-points split by drug and volatility condition. No significant ketamine-related differences in reaction times were observed for either valence or arousal. Black dots and error bars denote group means ± within-subject s.e.m. (Cousineau-Morey correction), coloured dots denote participant means. ** p < .01, * p < .05.

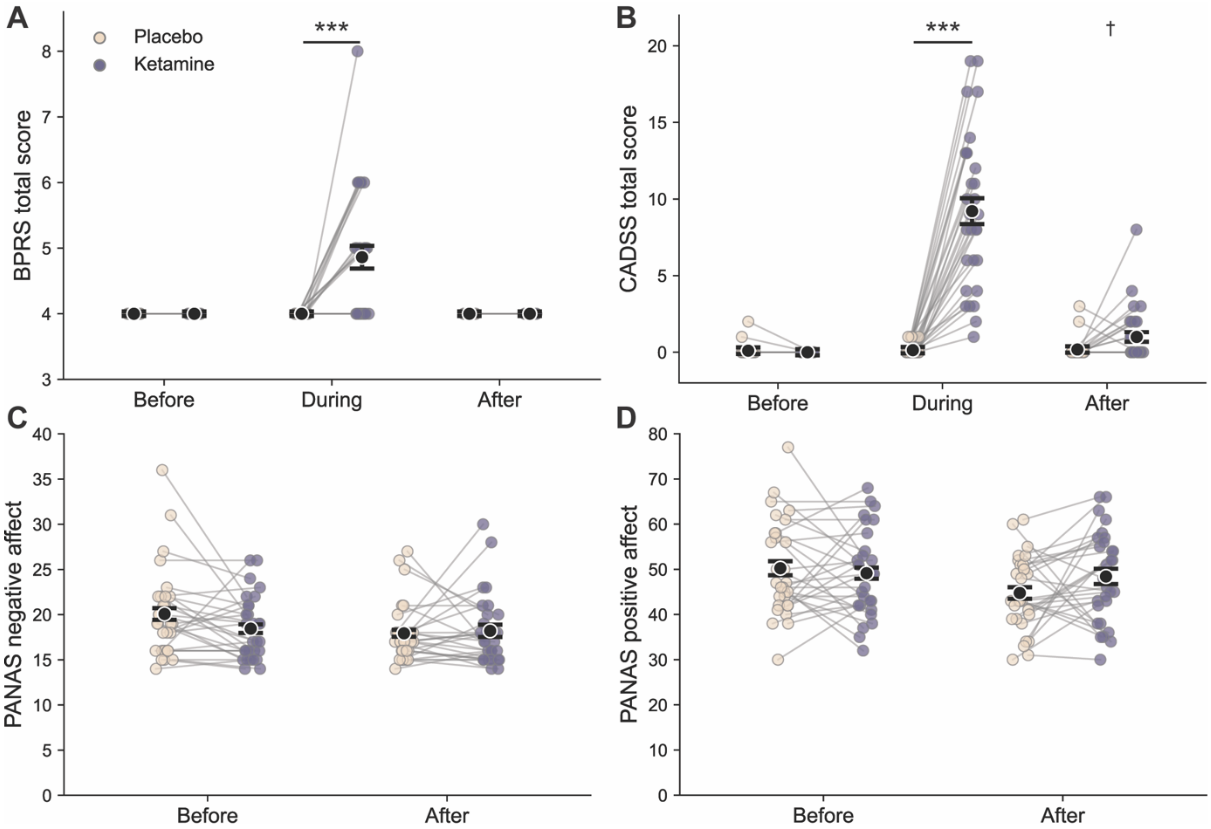


***Figure S6. Subjective effects of ketamine administration.*** **A.** Brief Psychiatric Rating Scale (BPRS) scores before, during, and after ketamine and placebo infusion. The post-infusion assessment took place immediately before the start of the continuous emotion rating task. Ketamine increased psychotomimetic symptoms during infusion, which had resolved by the post-infusion assessment. **B.** Clinician-Administered Dissociative States Scale (CADSS-6) scores before, during, and after infusion. Ketamine increased dissociative symptoms during infusion, which had largely subsided by the post-infusion assessment. **C.** Positive and Negative Affect Schedule (PANAS) negative affect scores before and after infusion. **D.** PANAS positive affect scores before and after infusion. Black dots and error bars denote group means ± within subject s.e.m. (Cousineau-Morey correction), coloured dots denote participant scores. BPRS and CADSS-6: ketamine vs placebo at each timepoint, paired Wilcoxon signed-rank tests, Holm-corrected *** $p < .001$, † $p < .10$. PANAS: linear mixed-effects models.

### Supplementary Results

#### *Replication of behavioural effects in the independent online sample*

The main behavioural effects were largely replicated in an independent online sample (n = 92; **Fig S1; Tables S5-8**). Online participants responded more rapidly than the in-person sample, with change-point detection beginning to increase approximately 0.5 s after change-point onset, and half of all change-points were detected within approximately 1.8s (**Fig S1A**). As in the in-person sample, change-point detection accuracy increased with change-point magnitude in both valence (b = 0.863, 95% CI [0.749, 0.976], $\chi^2(1) = 119.20$, $p < .001$) and arousal (b = 0.657, 95% CI [0.546, 0.768], $\chi^2(1) = 93.53$, $p < .001$; **Fig S1B**), while RTs decreased with increasing change-point magnitude (valence: b = -0.106, 95% CI [-0.124, -0.089], $F(1, 90.5) = 143.84$, $p < .001$; arousal: b = -0.174, 95% CI [-0.188, -0.161], $F(1, 9618.1) = 629.29$, $p < .001$; **Fig S1C**).

Emotion-dependent asymmetries in continuous tracking were also replicated. Rating errors were higher for positive than negative valence (b = 0.176, 95% CI [0.101, 0.251], F(1, 91) = 21.86, p < .001) and higher for low than high arousal (b = 0.507, 95% CI [0.397, 0.617], F(1, 91) = 83.63, p < .001; **Fig S1D**). At intermediate emotion levels, medium arousal showed a positive signed error (b = 0.492, 95% CI [0.423, 0.561], F(1, 91) = 201.29, p < .001), consistent with the in-person sample. In contrast to the small positive shift observed for neutral valence in the in-person sample, online participants showed a small negative signed error (b = -0.064, 95% CI [-0.113, -0.015], F(1, 91) = 6.77, p = .011), indicating a slight shift towards more negative ratings. The effects of change direction on RT also replicated in the online sample. Upward changes were slower than downward changes in valence (b = 0.087, 95% CI [0.062, 0.112], F(1, 90.2) = 47.45, p < .001), whereas upward changes were faster than downward changes in arousal (b = -0.231, 95% CI [-0.262, -0.200], F(1, 95) = 216.69, p < .001; **Fig S1F**). When examining the emotion level reached following a change, the arousal pattern was also consistent with the in-person sample: responses to medium arousal were slower than to low arousal following downward changes (b = 0.064, 95% CI [0.017, 0.112], F(1, 89.9) = 7.27, p = .008), and responses to high arousal were faster than to medium arousal following upward changes (b = -0.309, 95% CI [-0.371, -0.248], F(1, 89.8) = 99.42, p < .001; **Fig. S1G**). Valence showed some differences from the in-person sample, with faster responses to neutral than negative valence following downward changes (b = -0.102, 95% CI [-0.160, -0.043], F(1, 111) = 11.87, p < .001), while RTs did not differ between positive and neutral targets following upward changes (b = -0.029, 95% CI [-0.083, 0.025], F(1, 97.7) = 1.15, p = .287).

Finally, adaptation to environmental volatility partly replicated in the online cohort. Participants responded faster in volatile than stable blocks when rating valence (b = -0.131, 95% CI [-0.159, -0.103], F(1, 91) = 88.35, p < .001), whereas for arousal the difference did not reach significance (b = -0.020, 95% CI [-0.045, 0.005], F(1, 9613.1) = 2.44, p = .119; **Fig S1I**), as in the in-person sample. Change-point accuracy was lower in volatile blocks after accounting for change-point magnitude and change-point direction for valence (b = -0.272, 95% CI [-0.388, -0.155], $\chi^2(1)$ = 17.99, p < .001), but not significantly in arousal (b = -0.114, 95% CI [-0.222, -0.006], $\chi^2(1)$ = 3.84, p = .050). Face-wise integration kernels showed the same overall recency-weighting (**Fig S1J**), with steeper kernels in volatile blocks for valence (b = 0.038, 95% CI [0.013, 0.062], F(1, 91) = 9.02, p = .003, **Fig S1K**) but not arousal (b = -0.005, 95% CI [-0.022, 0.012], F(1, 91) = 0.36, p = .550). Together, these findings indicate that the principal effects of change magnitude, emotional context, and environmental volatility observed in the in-person cohort were largely reproduced in the independent online sample.

##### *Additional effects of environmental volatility*

In addition to the effects on reaction times and temporal evidence integration reported in the main text, environmental volatility influenced several measures of task performance (**Fig S3**). Participants made more rating changes in volatile than stable blocks in both valence (main effect of volatility: b = 0.276, 95% CI [0.237, 0.316], $\chi^2(1)$ = 59.12, p < .001) and arousal (b = 0.269, 95% CI [0.231, 0.307], $\chi^2(1)$ = 191.70, p < .001; **Fig S3A**). Relative to the number of true change-points, however, participants made proportionally more rating changes in stable than volatile blocks (**Fig S3B**). Change-point detection accuracy was lower in volatile than stable blocks after accounting for change-point magnitude and change-point direction in both valence (b = -0.311, 95% CI [-0.495, -0.128], $\chi^2(1)$ = 9.69, p = .002) and arousal (b = -0.295, 95% CI [-0.428, -0.162], $\chi^2(1)$ = 14.70, p < .001; **Fig S3C**). Together with the faster valence RTs in volatile blocks reported in the main text, these results indicate that participants adjusted their behaviour to environmental volatility, responding more frequently and rapidly when the underlying emotional state changed more often, although individual change-points were detected less accurately.

#### *Subjective effects of ketamine*

Ketamine produced transient psychotomimetic and dissociative effects during infusion that had largely resolved by the post-infusion assessment (**Fig S6A-B**). BPRS scores were at floor in all participants before and after infusion in both sessions and were higher during ketamine than placebo infusion (M = 4.86 vs 4.00; 15 of 29 participants higher on ketamine, none on placebo; $z = 3.46$, $p < .001$, $r = .64$). CADSS-6 scores were higher during ketamine than placebo in all 29 participants (M = 9.21 vs 0.14; $z = 4.69$, $p < .001$, $r = .87$) and did not differ before infusion (M = 0.00 vs 0.10; $z = 0.89$, $p = .371$, $r = .17$). After infusion, CADSS-6 scores no longer differed significantly between ketamine and placebo (M = 1.04 vs 0.18; $z = 2.19$, $p = .057$, $r = .41$), although some participants (10 of 28) still reported mild dissociative symptoms following ketamine (maximum score 8 of 24).

After infusion, PANAS scores did not differ between ketamine and placebo (**Fig S6C-D**). The drug x timepoint interaction was significant for negative affect (b = 1.897, 95% CI [0.123, 3.671], $F(1,56) = 4.59$, $p = .037$). This was driven by higher negative affect before placebo than before the ketamine infusion (b = -1.47, 95% CI [-3.155, 0.213], $t(48.48) = -1.76$, Holm p = .171), with similar levels after infusion (b = 0.43, 95% CI [-1.258, 2.109], $t(48.48) = 0.51$, Holm p = .614). The interaction was not significant for PANAS positive affect (b = 4.76, 95% CI [-0.52, 10.03], $F(1,56) = 3.264$, $p = .076$). Thus, participants performed the behavioural task after the marked acute psychotomimetic and dissociative effects of ketamine had subsided, without clear evidence for post-infusion changes in positive or negative affect.

BPRS and CADSS scores were at or near floor in most conditions. We therefore compared ketamine and placebo at each timepoint using paired Wilcoxon signed-rank tests, Holm-corrected across the three timepoints, with effect size $r = z/\sqrt{N}$. PANAS scores were analysed using linear mixed-effects models with drug, timepoint, their interaction and visit as fixed effects and random intercepts for participants and for each of their two infusion visits.

Treatment allocation was assessed retrospectively in a debrief questionnaire administered at the end of the study. Debrief data were available for 23 of 29 participants, all of whom correctly identified which visit involved ketamine and reported high confidence in their judgement.

#### *Deviations from preregistration*

The preregistration specified a logistic regression predicting whether a rating update followed each face, with the mean prediction error in five 0.5s time bins before the face as predictors. During model development, after unblinding but before examining drug effects, we identified two properties of this design that limited its interpretability. First, bin width is a free parameter, and the effect of volatility on integration-kernel steepness depended strongly on it. Second, adjacent time bins were highly correlated (mean $r \approx 0.79$ at 30 frames), increasingly so at narrower widths. We therefore changed to a face-wise linear kernel as the primary analysis (see Methods). Its regressors correspond to individual faces, which removes the bin-width parameter, and its outcome is the signed size of each update rather than whether an update occurred or not. We used linear rather than logistic regression because the task elicits graded updates. Changes in the emotional underlying true mean differed in magnitude and participants expressed larger updates through several consecutive rating steps. A binary outcome discards both the direction and the size of an update, whereas a linear model of the signed update (-2 to +2 steps) uses this information, and its weights are expressed directly in rating units per unit of prediction error. The time-binned kernel is reported in **Fig S4** and **Supplementary Table S3b**. With this kernel, the valence drug x volatility interaction effect on IKS observed with the face-wise kernel was not significant (p = .223), and no drug effect on IKS or BW-2 reached significance.

The preregistered random-effects structure (random slopes for drug, volatility and their interaction) could not be estimated for kernel measures, which have one value per participant and condition. We therefore used a sequence of simplifications, dropping random slopes while the fit was singular. The structures reached are reported in the supplementary tables.

#### Supplementary Tables

For each outcome, the main text reports the model we considered most appropriate for the effect of interest, typically the simplest model containing that effect and its random slope. The tables below additionally report alternative models that could have been used, including models with all predictors entered together and sensitivity models with additional covariates, so that readers can assess the robustness of the results. The direction and significance of the effects were the same across models.

##### *Behavioural results for the in-person cohort, n=29*

| **Predictor** | ***b*** | **95% CI** | | ***df*** | **F (KR)** | ***p*** |
|---|---|---|---|---|---|---|
| **DV: Signed RE (neutral in valence or medium in arousal)** | | | | | | |
| Neutral valence signed RE ~ drug + (1 \| subject) | | | | | | |
| Intercept | 0.059 | -0.002 | 0.12 | 1, 28 | 3.893 | .058 |
| Drug: ketamine - placebo | -0.021 | -0.111 | 0.069 | 1, 318 | 0.213 | .645 |
| Medium arousal Signed RE ~ drug + (1 + drug \| subject) | | | | | | |
| Intercept | 0.469 | 0.356 | 0.582 | 1, 28 | 72.534 | < .001 |
| Drug: ketamine - placebo | 0.115 | 0.015 | 0.216 | 1, 28 | 5.561 | .026 |
| **DV: valence signed RE flipped for positive emotion level; restricted to positive and negative emotion levels** | | | | | | |
| Valence RE ~ drug * emotion level + (1 + emotion level \| subject) | | | | | | |
| Intercept | 0.891 | 0.845 | 0.938 | 1, 28 | 1532.969 | < .001 |
| Drug: ketamine-placebo | -0.012 | -0.055 | 0.031 | 1, 636 | 0.284 | .594 |
| Emotion level: positive – negative | 0.132 | 0.027 | 0.237 | 1, 28 | 6.595 | .016 |
| Drug : Emotion level | 0.118 | 0.032 | 0.204 | 1, 636 | 7.216 | .007 |
| *Negative: ketamine – placebo* | *-0.071* | -0.140 | -0.001 | *636* | *-2.277* | .046 |
| *Positive: ketamine – placebo* | *0.047* | -0.022 | 0.117 | *636* | *1.522* | .128 |
| **DV: arousal signed RE flipped for high emotion level; restricted to high and low emotion levels** | | | | | | |
| Arousal RE ~ drug * emotion level + (1 + emotion level \| subject) | | | | | | |
| Intercept | 1.343 | 1.268 | 1.419 | 1, 28 | 1333.548 | < .001 |
| Drug: ketamine-placebo | -0.021 | -0.083 | 0.04 | 1, 636 | 0.462 | .497 |
| Emotion level: high – low | -0.453 | -0.643 | -0.264 | 1, 28 | 23.977 | < .001 |
| Drug : Emotion level | -0.123 | -0.246 | 0.001 | 1, 636 | 3.82 | .051 |

*Overview results from mixed-effects linear regression models*. Framewise rating errors (rating – true mean) were averaged within participant x drug x block x emotion level, such that each observation represented the mean rating error for one emotion level within a block. For the extreme emotion levels, errors were sign-flipped at the positive valence and high arousal pole. Neutral valence and medium arousal RE are unflipped, so positive values indicate ratings above the true mean. Mixed-effects linear regression models were fit to participants' rating errors with the lmer() function from lme4 package in R and tested with Kenward-Roger F tests. All factors were coded with sum-to-zero coding (emotion level: negative/low = -0.5, positive/high = 0.5; drug: placebo = -0.5, ketamine = 0.5) such that regression coefficients can be interpreted as the second level minus the first. Where the maximal model produced singular fits, unsupported random slopes/correlations were removed, and the selected structure is reported in the corresponding model line. The significant valence drug x emotion interaction was followed by simple effects of drug within each emotion level (Holm-corrected across the two levels, Bonferroni adjusted CIs)

**Table S1** Mixed-effects analyses of continuous rating errors across emotion levels and drug conditions.

| **Predictor** | ***b*** | **95% CI** | | ***df*** | **F/t/$\chi^2$** | ***p*** |
|---|---|---|---|---|---|---|
| **DV: reaction times (RT)** | | | | | | |
| logRT ~ dimension + (1 + dimension \| subject) | | | | | | |
| Intercept | 0.825 | 0.797 | 0.853 | 1, 28.0 | 3693.597 | < .001 |
| Dimension: arousal - valence | 0.011 | -0.017 | 0.039 | 1, 27.9 | 0.630 | .434 |
| Valence logRT ~ CP magnitude + (1 + CP magnitude \| subject) | | | | | | |
| Intercept | 0.809 | 0.776 | 0.843 | 1, 28.0 | 2463.888 | < .001 |
| Change point magnitude | -0.11 | -0.13 | -0.09 | 1, 27.9 | 125.177 | < .001 |
| Arousal logRT ~ CP magnitude + (1 + CP magnitude \|\| subject) | | | | | | |
| Intercept | 0.778 | 0.747 | 0.81 | 1, 29.6 | 2557.003 | < .001 |
| Change point magnitude | -0.136 | -0.152 | -0.12 | 1, 29.2 | 315.856 | < .001 |
| Valence logRT ~ CP direction + CP magnitude + (1 + CP direction \| subject) | | | | | | |
| Intercept | 0.809 | 0.776 | 0.842 | 1, 28.1 | 2509.277 | < .001 |
| CP direction: up - down | 0.091 | 0.063 | 0.12 | 1, 27.9 | 43.814 | < .001 |
| Change point magnitude | -0.108 | -0.123 | -0.094 | 1, 6076.7 | 209.122 | < .001 |
| Arousal logRT ~ CP direction + CP magnitude + (1 + CP direction \| subject) | | | | | | |
| Intercept | 0.779 | 0.747 | 0.811 | 1, 29.8 | 2511.842 | < .001 |
| CP direction: up - down | -0.117 | -0.148 | -0.086 | 1, 29.8 | 59.371 | < .001 |
| Change point magnitude | -0.122 | -0.137 | -0.108 | 1, 5473.3 | 278.006 | < .001 |
| Valence logRT ~ emotion level + CP magnitude + (1 + emotion level \| subject) – downward CP; excluding large CP magnitudes | | | | | | |
| Intercept | 0.809 | 0.768 | 0.85 | 1, 50.0 | 1544.997 | < .001 |
| Emotion level: neutral – negative | -0.004 | -0.059 | 0.05 | 1, 27.9 | 0.029 | .867 |
| CP magnitude (small and medium) | -0.006 | -0.051 | 0.038 | 1, 1666.6 | 0.074 | .785 |
| *Valence logRT ~ emotion level + CP magnitude + (1 + emotion level \| subject) – downward CP* | | | | | | |
| *Intercept* | 0.766 | 0.727 | 0.805 | 1, 28.1 | 1591.909 | < .001 |
| *Emotion level: neutral – negative* | 0.024 | -0.025 | 0.073 | 1, 31.1 | 0.983 | .329 |
| *CP magnitude* | -0.084 | -0.105 | -0.062 | 1, 2647.0 | 58.838 | < .001 |
| Valence logRT ~ emotion level + CP magnitude + (1 + emotion level \| subject) – upward CP; excluding large CP magnitudes | | | | | | |
| Intercept | 0.751 | 0.710 | 0.792 | 1, 55.8 | 1354.781 | < .001 |
| Emotion level: positive – neutral | -0.011 | -0.073 | 0.050 | 1, 27.6 | 0.143 | .709 |
| CP magnitude | -0.249 | -0.294 | -0.204 | 1, 1815.3 | 117.050 | < .001 |
| *Valence logRT ~ emotion level + CP magnitude + (1 + emotion level \| subject) – upward CP* | | | | | | |
| *Intercept* | 0.839 | 0.802 | 0.877 | 1, 29.0 | 2105.800 | < .001 |
| *Emotion level: positive – neutral* | 0.006 | -0.058 | 0.071 | 1, 31.4 | 0.037 | .848 |
| *CP magnitude* | -0.106 | -0.131 | -0.081 | 1, 2753.5 | 68.677 | < .001 |

| | | | | | | |
|---|---|---|---|---|---|---|
| Arousal logRT ~ emotion level + (1 + emotion level \| subject) – downward CP; excluding large CP magnitudes | | | | | | |
| Intercept | 0.955 | 0.928 | 0.982 | 1, 27.6 | 5255.452 | < .001 |
| Emotion level: medium – low | 0.086 | 0.036 | 0.136 | 1, 27.5 | 12.266 | .002 |
| *Arousal logRT ~ emotion level + CP magnitude + (1 + emotion level \| subject) – downward CP* | | | | | | |
| *Intercept* | 0.857 | 0.823 | 0.890 | 1, 45.3 | 2703.045 | < .001 |
| *Emotion level: medium – low* | 0.087 | 0.042 | 0.132 | 1, 38.7 | 15.193 | < .001 |
| *CP magnitude* | -0.100 | -0.124 | -0.077 | 1, 2939.2 | 68.707 | < .001 |
| Arousal logRT ~ emotion level + (1 + emotion level \| subject) – upward CP; excluding large CP magnitudes | | | | | | |
| Intercept | 0.846 | 0.809 | 0.884 | 1, 27.7 | 2166.701 | < .001 |
| Emotion level: high – medium | -0.214 | -0.290 | -0.137 | 1, 27.7 | 32.457 | < .001 |
| *Arousal logRT ~ emotion level + CP magnitude + (1 + emotion level \| subject) – upward CP* | | | | | | |
| *Intercept* | 0.777 | 0.734 | 0.820 | 1, 31.2 | 1366.997 | < .001 |
| *Emotion level: high – medium* | -0.213 | -0.280 | -0.146 | 1, 39.2 | 41.138 | < .001 |
| *CP magnitude* | -0.072 | -0.094 | -0.049 | 1, 2503.9 | 39.265 | < .001 |
| Valence logRT ~ volatility x CP magnitude + CP direction + (1 + volatility + CP magnitude \|\| subject) | | | | | | |
| Intercept | 0.815 | 0.782 | 0.849 | 1, 29.2 | 2487.327 | < .001 |
| Volatility: volatile – stable | -0.068 | -0.101 | -0.034 | 1, 31.7 | 17.181 | < .001 |
| CP direction: up - down | 0.088 | 0.064 | 0.113 | 1, 6043.5 | 50.670 | < .001 |
| CP magnitude | | | | 2, 76.4 | 68.146 | < .001 |
| Volatility : CP magnitude | | | | 2, 6058.5 | 4.915 | .007 |
| *Small jumps: volatile - stable* | -0.119 | -0.173 | -0.065 | 108.2 | -5.399 | < .001 |
| *Medium jumps: volatile - stable* | -0.016 | -0.084 | 0.051 | 277.1 | -0.576 | .565 |
| *Large jumps: volatile - stable* | -0.068 | -0.129 | -0.008 | 173.6 | -2.733 | .014 |
| Arousal logRT ~ volatility x CP magnitude + CP direction + (1 + volatility + CP magnitude \|\| subject) | | | | | | |
| Intercept | 0.776 | 0.744 | 0.808 | 1, 30.3 | 2429.999 | < .001 |
| Volatility: volatile – stable | 0.020 | -0.021 | 0.060 | 1, 34.0 | 0.986 | .328 |
| CP direction: up – down | -0.120 | -0.146 | -0.093 | 1, 5458.5 | 77.895 | < .001 |
| CP magnitude | | | | 1, 36.7 | 232.483 | < .001 |
| Volatility : CP magnitude | | | | 1, 5463.5 | 0.001 | .979 |
| Valence logRT ~ drug x volatility + CP magnitude + CP direction + (1 + drug + volatility \| subject) | | | | | | |
| Intercept | 0.822 | 0.788 | 0.855 | 1, 28.1 | 2523.766 | < .001 |
| Drug: ketamine - placebo | 0.002 | -0.027 | 0.031 | 1, 32.0 | 0.017 | .898 |
| Volatility: volatile – stable | -0.084 | -0.116 | -0.051 | 1, 27.9 | 28.095 | < .001 |
| CP magnitude | -0.107 | -0.121 | -0.092 | 1, 6056.5 | 203.067 | < .001 |
| CP direction: up - down | 0.093 | 0.069 | 0.117 | 1, 6033.2 | 57.801 | < .001 |
| Drug : volatility | -0.006 | -0.057 | 0.044 | 1, 6052.9 | 0.065 | .799 |
| Arousal logRT ~ drug x volatility + CP magnitude + CP direction + (1 + drug + volatility \| subject) | | | | | | |
| Intercept | 0.776 | 0.744 | 0.808 | 1, 29.8 | 2416.798 | < .001 |
| Drug: ketamine - placebo | -0.014 | -0.049 | 0.021 | 1, 31.2 | 0.673 | .418 |
| Volatility: volatile – stable | 0.019 | -0.021 | 0.058 | 1, 27.9 | 0.955 | .337 |
| CP magnitude | -0.123 | -0.137 | -0.109 | 1, 5449.3 | 282.032 | < .001 |
| CP direction: up - down | -0.119 | -0.146 | -0.093 | 1, 5444.4 | 77.958 | < .001 |
| Drug : volatility | -0.007 | -0.061 | 0.047 | 1, 5448.6 | 0.069 | .793 |

*Overview results from mixed-effects linear regression models.* Reaction times were analysed at the change-point level, with one observation per correctly detected change-point with a valid response time; RTs were log-transformed before analysis. Mixed-effects linear regression models were fit to participants' reaction times (logRT) with the `lmer()` function from the `lme4` package in R. All predictors were coded with sum-to-zero coding (dimension: valence = -0.5, arousal = 0.5; CP direction: down = -0.5, up = 0.5; volatility: stable = -0.5, volatile = 0.5; drug: placebo = -0.5, ketamine = 0.5). CP magnitude was centred at dimension-specific midpoints (valence: magnitude - 2; arousal: magnitude - 3). In the exploratory volatility x CP magnitude models, CP magnitude was instead entered as a sum-coded categorical factor so that the volatility effect could be estimated at each magnitude. In these models the volatility row is the volatility effect averaged across magnitudes, and the magnitude and interaction terms are multi-df F tests. Fixed effects were tested with Kenward-Roger F tests (lmerTest). Where the volatility x CP magnitude interaction was significant, volatile-stable contrasts were computed within each magnitude (emmeans, Kenward-Roger df), Holm-corrected across magnitudes, with Bonferroni-adjusted 95% CIs. Where the maximal random-effects structure produced singular fits, unsupported random slopes/correlations were removed and the selected structure is reported in the corresponding model line. Control analyses are shown in italics.

**Table S2.** Mixed-effects analyses of reaction times to detected change-points.

| **Predictor** | ***b*** | **95% CI** | | ***df*** | **F (KR) / t** | ***p*** |
|---|---|---|---|---|---|---|
| **DV: Integration kernel steepness (IKS) for the face-wise linear regression-based kernels** | | | | | | |
| Valence IKS ~ drug x volatility + (1 \| subject) | | | | | | |
| Intercept | -0.022 | -0.038 | -0.006 | 1, 28.0 | 7.925 | .009 |
| Drug: ketamine - placebo | 0.004 | -0.027 | 0.035 | 1, 84.0 | 0.061 | .806 |
| Volatility: volatile - stable | 0.028 | -0.003 | 0.059 | 1, 84.0 | 3.241 | .075 |
| Drug : volatility | 0.077 | 0.015 | 0.139 | 1, 84.0 | 6.030 | .016 |
| *Stable: ketamine - placebo* | -0.035 | -0.085 | 0.016 | 84.0 | -1.562 | .122 |
| *Volatile: ketamine - placebo* | 0.042 | -0.008 | 0.093 | 84.0 | 1.911 | .119 |
| *Ketamine: volatile-stable* | 0.067 | 0.016 | 0.117 | 84.0 | 3.009 | .007 |
| *Placebo: volatile-stable* | -0.010 | -0.061 | 0.040 | 84.0 | -0.463 | .644 |
| Arousal IKS ~ drug x volatility + (1 \| subject) | | | | | | |
| Intercept | 0.024 | 0.009 | 0.039 | 1, 28.0 | 11.258 | .002 |
| Drug: ketamine - placebo | 0.006 | -0.018 | 0.03 | 1, 84.0 | 0.245 | .622 |
| Volatility: volatile - stable | -0.019 | -0.043 | 0.005 | 1, 84.0 | 2.416 | .124 |
| Drug : volatility | 0.001 | -0.047 | 0.05 | 1, 84.0 | 0.003 | .96 |
| **DV: beta weight of the second most recent regressor ($BW_{-2}$), stable blocks** | | | | | | |
| Valence $BW_{-2}$: ketamine - placebo | 0.019 | -0.014 | 0.051 | 28 | 1.178 | .249 |
| Arousal $BW_{-2}$: ketamine - placebo | -0.003 | -0.027 | 0.02 | 28 | -0.295 | .770 |

*Overview of mixed-effects linear regression models.* Integration kernel steepness (IKS) was averaged across blocks within participant x drug x dimension x volatility condition before analysis, such that each observation represented one participant's mean IKS for a given drug and volatility condition. Models were fit using the lmer() function from the lme4 package in R. Drug and volatility were coded using sum-to-zero contrasts (placebo = -0.5, ketamine = 0.5; stable = -0.5, volatile = 0.5), such that coefficients represent the second level minus the first. Where the maximal random-effects structure produced singular fits, unsupported random slopes/correlations were removed, and the selected structure is reported in the corresponding model line. For valence, the significant drug x volatility interaction was followed by pairwise simple effects of drug (ketamine - placebo) within each volatility level, and of volatility within each drug condition, Holm-corrected within each pair; these follow-ups are reported as t ratios. Drug effect on stable block BW-2 was tested with the preregistered two-sided paired t-tests.

**Table S3.** Mixed-effects analyses of face-wise linear regression-based integration kernel steepness across drug and volatility conditions.

| **Predictor** | ***b*** | **95% CI** | | ***df*** | **F (KR) / t** | ***p*** |
|---|---|---|---|---|---|---|
| **DV: Integration kernel steepness (IKS) for the 30-frame time-binned logistic regression-based kernels** | | | | | | |
| Valence IKS ~ drug x volatility + (1 + volatility \| subject) | | | | | | |
| Intercept | -0.101 | -0.180 | -0.022 | 1, 28.0 | 6.866 | .014 |
| Drug: ketamine - placebo | -0.085 | -0.203 | 0.032 | 1, 56.0 | 2.101 | .153 |
| Volatility: volatile - stable | 0.128 | -0.035 | 0.290 | 1, 28.0 | 2.597 | .118 |
| Drug : volatility | 0.145 | -0.091 | 0.380 | 1, 56.0 | 1.516 | .223 |
| Arousal IKS ~ drug x volatility + (1 + volatility \| subject) | | | | | | |

| | | | | | | |
|---|---|---|---|---|---|---|
| Intercept | 0.194 | 0.067 | 0.320 | 1, 28.0 | 9.813 | .004 |
| Drug: ketamine - placebo | 0.163 | -0.026 | 0.352 | 1, 56.0 | 2.995 | .089 |
| Volatility: volatile - stable | 0.030 | -0.222 | 0.281 | 1, 28.0 | 0.058 | .812 |
| Drug : volatility | 0.066 | -0.312 | 0.443 | 1, 56.0 | 0.121 | .729 |
| **DV: beta weight of the second most recent time bin ($BW_{-2}$), stable blocks** | | | | | | |
| Valence $BW_{-2}$: ketamine - placebo | 0.074 | -0.050 | 0.198 | 28.0 | 1.227 | .230 |
| Arousal $BW_{-2}$: ketamine - placebo | -0.097 | -0.336 | 0.142 | 28.0 | -0.831 | .413 |

*Overview of mixed-effects linear regression models.* Integration kernel steepness (IKS) was averaged across blocks within participant x drug x dimension x volatility condition before analysis, such that each observation represented one participant's mean IKS for a given drug and volatility condition. Models were fit using the lmer() function from the lme4 package in R. Drug and volatility were coded using sum-to-zero contrasts (placebo = -0.5, ketamine = 0.5; stable = -0.5, volatile = 0.5), such that coefficients represent the second level minus the first. Where the preregistered maximal random-effects structure produced singular fits, unsupported random slopes/correlations were removed, and the selected structure is reported in the corresponding model line.

**Table S3b.** Mixed-effects analyses of preregistered 30-frame time-binned logistic regression-based integration kernel steepness across drug and volatility conditions.

| **Predictor** | ***b*** | **95% CI** | | ***df*** | **$z/\chi^2$** | ***p*** |
|---|---|---|---|---|---|---|
| **DV: change-point accuracy** | | | | | | |
| Valence CP Accuracy ~ CP magnitude + (1+ CP magnitude \| subject) | | | | | | |
| Intercept | 1.564 | 1.356 | 1.772 | | 14.74 | < .001 |
| Change point magnitude | 0.888 | 0.763 | 1.014 | 1 | 61.493 | < .001 |
| Arousal CP Accuracy ~ CP magnitude + (1+ CP magnitude \| subject) | | | | | | |
| Intercept | 1.239 | 1.028 | 1.451 | | 11.471 | < .001 |
| Change point magnitude | 0.61 | 0.511 | 0.709 | 1 | 55.598 | < .001 |
| Valence CP Accuracy ~ CP direction + CP magnitude + (1 + CP direction \|\| subject) | | | | | | |
| Intercept | 1.512 | 1.34 | 1.685 | | 17.155 | < .001 |
| CP direction: up - down | 0.16 | 0.048 | 0.273 | 1 | 7.615 | .006 |
| Change point magnitude | 0.816 | 0.739 | 0.893 | 1 | 503.021 | < .001 |
| Arousal CP Accuracy ~ CP direction + CP magnitude + (1 \| subject) | | | | | | |
| Intercept | 1.199 | 1.02 | 1.378 | | 13.141 | < .001 |
| CP direction: up - down | -0.168 | -0.27 | -0.066 | 1 | 10.436 | .001 |
| Change point magnitude | 0.58 | 0.512 | 0.649 | 1 | 321.744 | < .001 |
| Valence CP Accuracy ~ emotion level + CP magnitude + (1 + emotion level \| subject) – downward CP; excluding large CP magnitudes | | | | | | |
| Intercept | 1.554 | 1.353 | 1.756 | | 15.089 | < .001 |
| Emotion level: neutral – negative | 0.033 | -0.222 | 0.288 | 1 | 0.064 | .801 |
| CP magnitude: medium – small | 1.108 | 0.914 | 1.302 | 1 | 134.853 | < .001 |
| Valence CP Accuracy ~ emotion level + CP magnitude + (1 + emotion level \| subject) – upward CP; excluding large CP magnitudes | | | | | | |
| Intercept | 1.579 | 1.339 | 1.82 | | 12.865 | < .001 |
| Emotion level: positive – neutral | -0.197 | -0.43 | 0.035 | 1 | 2.769 | .096 |
| CP magnitude: medium – small | 0.849 | 0.666 | 1.032 | 1 | 84.668 | < .001 |
| Arousal CP Accuracy ~ emotion level + (1 + emotion level \| subject) – downward CP; excluding large CP magnitudes | | | | | | |
| Intercept | 0.722 | 0.54 | 0.904 | | 7.789 | < .001 |
| Emotion level: medium – low | -0.318 | -0.565 | -0.071 | 1 | 5.777 | .016 |
| Arousal CP Accuracy ~ emotion level + (1 + emotion level \| subject) – upward CP; excluding large CP magnitudes | | | | | | |
| Intercept | 0.543 | 0.376 | 0.71 | | 6.374 | < .001 |
| Emotion level: high – medium | -0.075 | -0.35 | 0.2 | 1 | 0.285 | .594 |
| Valence CP Accuracy ~ drug x volatility + CP magnitude + CP direction + (1 + drug x volatility \| subject) | | | | | | |

| | | | | | | |
|---|---|---|---|---|---|---|
| Intercept | 1.587 | 1.407 | 1.767 | | 17.259 | < .001 |
| Drug: ketamine - placebo | -0.086 | -0.251 | 0.08 | 1 | 1.025 | .311 |
| Volatility: volatile – stable | -0.311 | -0.495 | -0.128 | 1 | 9.687 | .002 |
| CP magnitude | 0.838 | 0.76 | 0.916 | 1 | 514.967 | < .001 |
| CP direction: up - down | 0.148 | 0.038 | 0.257 | 1 | 6.924 | .009 |
| Drug : volatility | 0.16 | -0.093 | 0.414 | 1 | 1.508 | .219 |
| Arousal CP Accuracy ~ drug x volatility + CP magnitude + CP direction + (1 + drug + volatility \| subject) | | | | | | |
| Intercept | 1.262 | 1.086 | 1.438 | | 14.059 | < .001 |
| Drug: ketamine - placebo | -0.041 | -0.189 | 0.107 | 1 | 0.29 | .590 |
| Volatility: volatile – stable | -0.295 | -0.428 | -0.162 | 1 | 14.698 | < .001 |
| CP magnitude | 0.588 | 0.519 | 0.657 | 1 | 325.987 | < .001 |
| CP direction: up - down | -0.161 | -0.264 | -0.059 | 1 | 9.499 | .002 |
| Drug : volatility | -0.299 | -0.515 | -0.082 | 1 | 7.288 | .007 |
| *Stable: ketamine - placebo* | 0.017 | -0.02 | 0.054 | | 1.041 | .298 |
| *Volatile: ketamine - placebo* | -0.035 | -0.07 | -0.001 | | -2.286 | .045 |
| *Ketamine: volatile – stable* | -0.077 | -0.113 | -0.042 | | -4.844 | < .001 |
| *Placebo: volatile – stable* | -0.025 | -0.058 | 0.009 | | -1.667 | .095 |

*Overview results from mixed-effects logistic regression models*. Change-point detection accuracy was analysed at the individual change-point level, with one binary observation per change-point (0 = miss, 1 = hit). Models were fit using the glmer() function from the lme4 package in R (binomial family, logit link). Binary predictors were coded with -0.5/+0.5 sum-to-zero contrasts (CP direction: down = -0.5, up = +0.5; volatility: stable = -0.5, volatile = +0.5; drug: placebo = -0.5, ketamine = +0.5), so that coefficients reflect the second level minus the first on the logit (log-odds) scale. CP magnitude was centred at dimension-specific midpoints (valence: magnitude - 2; arousal: magnitude - 3). Where the maximal random-effects structure produced singular fits, unsupported random slopes/correlations were removed and the selected structure is reported in the corresponding model line. Fixed effects (b, log-odds scale) were tested with likelihood-ratio $\chi^2$ tests comparing the full model with a model without that term; intercepts are reported with Wald z. The significant arousal drug x volatility interaction was followed by simple effects of drug within each volatility level and of volatility within each drug condition, expressed as differences in hit probability (emmeans, response scale; z tests, Holm-corrected within each pair of contrasts).

**Table S4.** Mixed-effects analyses of change-point detection accuracy across task and drug conditions.

#### *Behavioural results for the online cohort, n=92*

| **Predictor** | ***b*** | **95% CI** | | ***df*** | **F (KR)** | ***p*** |
|---|---|---|---|---|---|---|
| **DV: valence signed rating error (RE) flipped for positive emotion level; restricted to positive and negative emotion levels** | | | | | | |
| RE ~ emotion level + (1 + emotion level \| participant) | | | | | | |
| Intercept | 0.848 | 0.811 | 0.885 | 1, 91.0 | 2126.937 | < .001 |
| Emotion level: positive – negative | 0.176 | 0.101 | 0.251 | 1, 91.0 | 21.859 | < .001 |
| **DV: arousal signed RE flipped for high emotion level; restricted to high and low emotion levels** | | | | | | |
| RE ~ emotion level + (1 + emotion level \| participant) | | | | | | |
| Intercept | 1.24 | 1.175 | 1.306 | 1, 91.0 | 1421.01 | < .001 |
| Emotion level: high – low | -0.507 | -0.617 | -0.397 | 1, 91.0 | 83.631 | < .001 |
| **DV: Signed RE (neutral in valence or medium in arousal)** | | | | | | |
| Signed RE ~ 1 + (1 \| participant) fit separately for valence and arousal | | | | | | |
| Intercept (neutral valence) | -0.064 | -0.113 | -0.015 | 1, 91.0 | 6.768 | .011 |
| Intercept (medium arousal) | 0.492 | 0.423 | 0.561 | 1, 91.0 | 201.29 | < .001 |

*Overview results from mixed-effects linear regression models*. As Table S1, for the online cohort (n=92), without drug terms. Neutral valence and medium arousal signed errors were tested against zero in separate intercept-only models.

**Table S5** Mixed-effects analyses of continuous rating errors across emotion levels in the online cohort (n=92)

| **Predictor** | ***b*** | **95% CI** | | ***df*** | **F/t/$\chi^2$** | ***p*** |
|---|---|---|---|---|---|---|
| **DV: reaction times (RT)** | | | | | | |
| logRT ~ dimension + (1 + dimension \| subject) | | | | | | |
| Intercept | 0.506 | 0.474 | 0.538 | 1, 91.0 | 968.41 | < .001 |
| Dimension: arousal - valence | 0.003 | -0.022 | 0.029 | 1, 90.7 | 0.074 | .787 |
| Valence logRT ~ CP magnitude + (1 + CP magnitude \| subject) | | | | | | |
| Intercept | 0.489 | 0.453 | 0.524 | 1, 91.0 | 743.072 | < .001 |
| Change point magnitude | -0.106 | -0.124 | -0.089 | 1, 90.5 | 143.841 | < .001 |
| Arousal logRT ~ CP magnitude + (1 \| subject) | | | | | | |
| Intercept | 0.429 | 0.394 | 0.464 | 1, 96.7 | 581.464 | < .001 |
| Change point magnitude | -0.174 | -0.188 | -0.161 | 1, 9618.1 | 629.29 | < .001 |
| Valence logRT ~ CP direction + CP magnitude + (1 + CP direction \| subject) | | | | | | |
| Intercept | 0.488 | 0.454 | 0.522 | 1, 91.6 | 793.558 | < .001 |
| CP direction: up - down | 0.087 | 0.062 | 0.112 | 1, 90.2 | 47.454 | < .001 |
| Change point magnitude | -0.108 | -0.123 | -0.094 | 1, 10409.3 | 219.037 | < .001 |
| Arousal logRT ~ CP direction + CP magnitude + (1 + CP direction \| subject) | | | | | | |
| Intercept | 0.431 | 0.395 | 0.467 | 1, 96.2 | 570.718 | < .001 |
| CP direction: up - down | -0.231 | -0.262 | -0.2 | 1, 95.0 | 216.691 | < .001 |
| Change point magnitude | -0.148 | -0.161 | -0.134 | 1, 9570.7 | 450.015 | < .001 |
| Valence logRT ~ emotion level + CP magnitude + (1 + emotion level \|\| subject) – downward CP; excluding large CP magnitudes | | | | | | |
| Intercept | 0.482 | 0.443 | 0.522 | 1, 223.3 | 585.717 | < .001 |
| Emotion level: neutral – negative | -0.102 | -0.16 | -0.043 | 1, 111.0 | 11.872 | < .001 |
| CP magnitude (small and medium) | -0.033 | -0.078 | 0.012 | 1, 2797.3 | 2.094 | .148 |
| Valence logRT ~ emotion level + CP magnitude + (1 + emotion level \|\| subject) – upward CP; excluding large CP magnitudes | | | | | | |
| Intercept | 0.424 | 0.382 | 0.466 | 1, 181.1 | 395.841 | < .001 |
| Emotion level: positive – neutral | -0.029 | -0.083 | 0.025 | 1, 97.7 | 1.145 | .287 |
| CP magnitude | -0.256 | -0.299 | -0.212 | 1, 3354.8 | 131.396 | < .001 |
| Arousal logRT ~ emotion level + (1 + emotion level \| subject) – downward CP; excluding large CP magnitudes | | | | | | |
| Intercept | 0.685 | 0.65 | 0.719 | 1, 90.7 | 1554.208 | < .001 |
| Emotion level: medium – low | 0.064 | 0.017 | 0.112 | 1, 89.9 | 7.27 | .008 |
| Arousal logRT ~ emotion level + (1 + emotion level \| subject) – upward CP; excluding large CP magnitudes | | | | | | |
| Intercept | 0.481 | 0.439 | 0.522 | 1, 90.6 | 536.922 | < .001 |
| Emotion level: high – medium | -0.309 | -0.371 | -0.248 | 1, 89.8 | 99.424 | < .001 |
| Valence logRT ~ volatility + CP magnitude + CP direction + (1 + volatility \| subject) | | | | | | |
| Intercept | 0.507 | 0.472 | 0.542 | 1, 91.8 | 830.176 | < .001 |
| Volatility: volatile – stable | -0.131 | -0.159 | -0.103 | 1, 91.0 | 88.348 | < .001 |
| CP magnitude | -0.104 | -0.118 | -0.089 | 1, 10422.3 | 201.688 | < .001 |
| CP direction: up - down | 0.087 | 0.063 | 0.110 | 1, 10366.6 | 52.113 | < .001 |
| Arousal logRT ~ volatility + CP magnitude + CP direction + (1 \| subject) | | | | | | |
| Intercept | 0.434 | 0.399 | 0.470 | 1, 98.9 | 581.824 | < .001 |
| Volatility: volatile – stable | -0.020 | -0.045 | 0.005 | 1, 9613.1 | 2.437 | .119 |
| CP magnitude | -0.148 | -0.162 | -0.134 | 1, 9615.4 | 450.778 | < .001 |
| CP direction: up - down | -0.234 | -0.258 | -0.209 | 1, 9613.2 | 347.490 | < .001 |

*Overview results from mixed-effects linear regression models.* As Table S2, for the online cohort (n = 92), without drug terms.

**Table S6.** Mixed-effects analyses of reaction times to detected change-points in the online cohort (n=92).

| Predictor | *b* | 95% CI | | *df* | F (KR) | *p* |
|---|---|---|---|---|---|---|
| **DV: Integration kernel steepness (IKS)** | | | | | | |
| Valence IKS ~ volatility + (1 \| subject) | | | | | | |
| Intercept | 0.159 | 0.141 | 0.177 | 1, 91.0 | 315.745 | < .001 |
| Volatility: volatile - stable | 0.038 | 0.013 | 0.062 | 1, 91.0 | 9.022 | .003 |
| Arousal IKS ~ volatility + (1 \| subject) | | | | | | |
| Intercept | 0.138 | 0.123 | 0.153 | 1, 91.0 | 343.663 | < .001 |
| Volatility: volatile - stable | -0.005 | -0.022 | 0.012 | 1, 91.0 | 0.360 | .550 |

*Overview of mixed-effects linear regression models. As Table S3, for the online cohort (n = 92) without the drug term.* Integration kernel steepness (IKS) was averaged across blocks within participant x dimension x volatility condition before analysis, such that each observation represented one participant's mean IKS for a given rating dimension and volatility condition.

**Table S7.** Mixed-effects analyses of integration kernel steepness across volatility conditions in the online cohort (n=92).

| Predictor | *b* | 95% CI | | *df* | F/t/$\chi^2$ | *p* |
|---|---|---|---|---|---|---|
| **DV: change-point accuracy** | | | | | | |
| Valence CP Accuracy ~ CP magnitude + (1 + CP magnitude \| subject) | | | | | | |
| Intercept | 2.072 | 1.905 | 2.239 | | 24.304 | < .001 |
| Change point magnitude | 0.863 | 0.749 | 0.976 | 1 | 119.204 | < .001 |
| Arousal CP Accuracy ~ CP magnitude + (1+ CP magnitude \| subject) | | | | | | |
| Intercept | 1.82 | 1.624 | 2.016 | | 18.215 | < .001 |
| Change point magnitude | 0.657 | 0.546 | 0.768 | 1 | 93.529 | < .001 |
| Valence CP Accuracy ~ CP direction + CP magnitude + (1 \| subject) | | | | | | |
| Intercept | 1.941 | 1.813 | 2.07 | | 29.631 | < .001 |
| CP direction: up - down | 0.309 | 0.211 | 0.406 | 1 | 38.534 | < .001 |
| Change point magnitude | 0.674 | 0.607 | 0.741 | 1 | 434.491 | < .001 |
| Arousal CP Accuracy ~ CP direction + CP magnitude + (1 + CP direction \| subject) | | | | | | |
| Intercept | 1.645 | 1.502 | 1.788 | | 22.544 | < .001 |
| CP direction: up - down | -0.191 | -0.298 | -0.083 | 1 | 10.933 | < .001 |
| Change point magnitude | 0.469 | 0.409 | 0.53 | 1 | 260.057 | < .001 |
| Valence CP Accuracy ~ emotion level + CP magnitude + (1 + emotion level \| subject) – downward CP; excluding large CP magnitudes | | | | | | |
| Intercept | 1.888 | 1.716 | 2.06 | | 21.524 | < .001 |
| Emotion level: neutral – negative | -0.244 | -0.449 | -0.039 | 1 | 5 | .025 |
| CP magnitude: medium – small | 1.122 | 0.949 | 1.295 | 1 | 177.682 | < .001 |
| Valence CP Accuracy ~ emotion level + CP magnitude + (1 + emotion level \| subject) – upward CP; excluding large CP magnitudes | | | | | | |
| Intercept | 2.043 | 1.866 | 2.22 | | 22.628 | < .001 |
| Emotion level: positive – neutral | -0.227 | -0.446 | -0.008 | 1 | 4.313 | .038 |
| CP magnitude: medium – small | 0.685 | 0.516 | 0.853 | 1 | 65.258 | < .001 |
| Arousal CP Accuracy ~ emotion level + (1 + emotion level \| subject) – downward CP; excluding large CP magnitudes | | | | | | |
| Intercept | 1.31 | 1.162 | 1.459 | | 17.321 | < .001 |
| Emotion level: medium – low | -0.132 | -0.326 | 0.062 | 1 | 1.76 | .185 |
| Arousal CP Accuracy ~ emotion level + (1 + emotion level \| subject) – upward CP; excluding large CP magnitudes | | | | | | |
| Intercept | 1.049 | 0.922 | 1.176 | | 16.187 | < .001 |
| Emotion level: high – medium | -0.123 | -0.302 | 0.056 | 1 | 1.792 | .181 |

| | | | | | | |
|---|---|---|---|---|---|---|
| Valence CP Accuracy ~ volatility + CP magnitude + CP direction + (1 + volatility \| subject) | | | | | | |
| Intercept | 1.997 | 1.868 | 2.125 | | 30.498 | < .001 |
| Volatility: volatile – stable | -0.272 | -0.388 | -0.155 | 1 | 17.987 | < .001 |
| CP magnitude | 0.691 | 0.623 | 0.76 | 1 | 448.538 | < .001 |
| CP direction: up - down | 0.29 | 0.192 | 0.388 | 1 | 33.624 | < .001 |
| Arousal CP Accuracy ~ volatility + CP magnitude + CP direction + (1 + volatility \| subject) | | | | | | |
| Intercept | 1.664 | 1.525 | 1.803 | | 23.532 | < .001 |
| Volatility: volatile – stable | -0.114 | -0.222 | -0.006 | 1 | 3.841 | .050 |
| CP magnitude | 0.473 | 0.412 | 0.534 | 1 | 263.153 | < .001 |
| CP direction: up - down | -0.17 | -0.26 | -0.08 | 1 | 13.526 | < .001 |

*Overview results from mixed-effects logistic regression models*. As for Table S4, for the online cohort (n = 92), without drug terms.

**Table S8.** Mixed-effects analyses of change-point detection accuracy across task conditions in the online cohort (n=92).

| **Predictor** | ***b*** | **95% CI** | | ***df*** | **F/t/$\chi^2$** | ***p*** |
|---|---|---|---|---|---|---|
| **DV: Integration kernel steepness (IKS) face-wise linear regression kernels (Table S3)** | | | | | | |
| Valence IKS ~ drug x volatility + visit order + gender + (1 \| subject) | | | | | | |
| Drug: ketamine - placebo | 0.004 | -0.027 | 0.035 | 1, 84.0 | 0.06 | .807 |
| Drug : volatility | 0.077 | 0.014 | 0.139 | 1, 84.0 | 5.995 | .016 |
| Arousal IKS ~ drug x volatility + visit order + gender + (1 \| subject) | | | | | | |
| Drug: ketamine - placebo | 0.006 | -0.018 | 0.03 | 1, 84.0 | 0.245 | .622 |
| Drug : volatility | 0.001 | -0.047 | 0.05 | 1, 84.0 | 0.003 | .96 |
| **DV: Integration kernel steepness (IKS) time-binned logistic regression kernels (Table S3b)** | | | | | | |
| Valence IKS ~ drug x volatility + visit order + gender + (1 + volatility \| subject) | | | | | | |
| Drug: ketamine - placebo | -0.085 | -0.203 | 0.032 | 1, 56.0 | 2.101 | .153 |
| Drug : volatility | 0.145 | -0.091 | 0.38 | 1, 56.0 | 1.516 | .223 |
| Arousal IKS ~ drug x volatility + visit order + gender + (1 + volatility \| subject) | | | | | | |
| Drug: ketamine - placebo | 0.163 | -0.026 | 0.352 | 1, 56.0 | 2.995 | .089 |
| Drug : volatility | 0.066 | -0.312 | 0.443 | 1, 56.0 | 0.121 | .729 |
| **DV: valence signed RE, positive and negative emotion levels (Table S1)** | | | | | | |
| Valence RE ~ drug x emotion level + visit order + gender + (1 + emotion level \| subject) | | | | | | |
| Drug: ketamine - placebo | -0.012 | -0.055 | 0.031 | 1, 636.0 | 0.284 | .594 |
| Drug : emotion level | 0.118 | 0.032 | 0.204 | 1, 636.0 | 7.216 | .007 |
| **DV: arousal signed RE, high and low emotion levels (Table S1)** | | | | | | |
| Arousal RE ~ drug x emotion level + visit order + gender + (1 + emotion level \| subject) | | | | | | |
| Drug: ketamine - placebo | -0.021 | -0.083 | 0.04 | 1, 636.0 | 0.462 | .497 |
| Drug : emotion level | -0.123 | -0.246 | 0.001 | 1, 636.0 | 3.82 | .051 |
| **DV: signed RE, neutral valence / medium arousal (Table S1)** | | | | | | |
| Neutral valence signed RE ~ drug + visit order + gender + (1 \| subject) | | | | | | |
| Drug: ketamine - placebo | -0.021 | -0.11 | 0.068 | 1, 318.0 | 0.213 | .645 |
| Medium arousal signed RE ~ drug + visit order + gender + (1 + drug \| subject) | | | | | | |
| Drug: ketamine - placebo | 0.115 | 0.019 | 0.211 | 1, 28.0 | 5.561 | .026 |
| **DV: reaction times (Table S2)** | | | | | | |
| Valence logRT ~ drug x volatility + CP magnitude + CP direction + visit order + gender + (1 + drug + volatility \| subject) | | | | | | |
| Drug: ketamine - placebo | 0.002 | -0.027 | 0.031 | 1, 32.0 | 0.02 | .888 |
| Drug : volatility | -0.006 | -0.056 | 0.044 | 1, 6051.9 | 0.057 | .812 |
| Arousal logRT ~ drug x volatility + CP magnitude + CP direction + visit order + gender + (1 + drug + volatility \| subject) | | | | | | |
| Drug: ketamine - placebo | -0.014 | -0.049 | 0.021 | 1, 31.2 | 0.665 | .421 |
| Drug : volatility | -0.007 | -0.061 | 0.047 | 1, 5448.7 | 0.066 | .797 |
| **DV: change-point accuracy (Table S4)** | | | | | | |

| | | | | | | |
|---|---|---|---|---|---|---|
| Valence CP accuracy ~ drug x volatility + CP magnitude + CP direction + visit order + gender + (1 + drug x volatility \| subject) | | | | | | |
| Drug: ketamine - placebo | -0.086 | -0.251 | 0.08 | 1 | 1.019 | .313 |
| Drug : volatility | 0.163 | -0.094 | 0.419 | 1 | 1.525 | .217 |
| Arousal CP accuracy ~ drug x volatility + CP magnitude + CP direction + visit order + gender + (1 + drug + volatility \| subject) | | | | | | |
| Drug: ketamine - placebo | -0.042 | -0.19 | 0.106 | 1 | 0.304 | .582 |
| Drug : volatility | -0.298 | -0.515 | -0.082 | 1 | 7.272 | .007 |
| **DV: number of rating changes** | | | | | | |
| Valence rating changes ~ drug x volatility + visit order + gender + (1 + drug x volatility \| subject) | | | | | | |
| Drug: ketamine - placebo | 0.022 | -0.038 | 0.083 | 1 | 0.52 | .471 |
| Drug : volatility | -0.002 | -0.075 | 0.071 | 1 | 0.003 | .959 |
| Arousal rating changes ~ drug x volatility + visit order + gender + (1 \| subject) | | | | | | |
| Drug: ketamine - placebo | -0.024 | -0.062 | 0.014 | 1 | 1.532 | .216 |
| Drug : volatility | -0.078 | -0.154 | -0.001 | 1 | 3.96 | .047 |

*Overview of the preregistered covariate sensitivity analyses*. All in-person drug models from Supplementary Tables S1-S4, and the rating-change models were refitted with visit order (ketamine first = 0.5, placebo first = -0.5) and gender (female = 0.5, male = -0.5) as additional fixed effects. Random-effects structures were identical to those of the corresponding main models. Only drug terms are shown. Linear mixed models were tested with Kenward-Roger F tests, and logistic (CP accuracy) and Poisson (rating changes) models with likelihood ratio $\chi2$ tests; b for these models is on the log-odds and log-rate scale, respectively. No drug effect changed significance.

**Table S9.** Drug effects with the preregistered covariates (visit order and gender).
